# SF3B1^K700E^ rewires splicing of cell-cycle regulators

**DOI:** 10.1101/2025.05.08.652831

**Authors:** Mai Baker, Eden Engal, Aveksha Sharma, Mayra Petasny, Shiri Jaffe-Herman, Ophir Geminder, Min-Hua Su, Mercedes Bentata, Gillian Kay, Maayan Salton

**Author notes:** These authors contributed equally (Mai Baker, Eden Engal, Aveksha Sharma, Mayra Petasny).

## Abstract

Pre-mRNA splicing plays a crucial role in maintaining cellular homeostasis, with strict regulation required for processes such as cell cycle progression. SF3B1, a core component of the spliceosome, has emerged as a key player in alternative splicing regulation and is frequently mutated in cancer. Among these mutations, SF3B1^K700E^ disrupts normal splicing patterns and deregulates cell cycle control. Here we profiled K562 erythroleukaemia cells expressing either wild-type or SF3B1^K700E^ by RNA-seq and uncovered 763 high-confidence splicing alterations enriched for G2/M regulators, including ARPP19, ENSA, STAG2 and ECT2. Notably, increased inclusion of ARPP19 exon 2 produces the ARPP19-long isoform, which sustains PP2A-B55 inhibition and promotes mitotic progression. A core subset of the K700E-linked splicing changes re-appeared after siRNA-mediated SF3B1 depletion in HeLa cells, underscoring a mutation-dependent spliceosomal signature that transcends cell type. Pharmacological inhibition of DYRK1A (INDY) or broad serine/threonine phosphatases (okadaic acid) shifted ARPP19 exon 2 inclusion in the same direction as SF3B1^K700E^, pointing to a kinase–phosphatase signaling axis that influences these splice events. Functionally, ectopic expression of ARPP19-long accelerated mitotic exit, and high ARPP19-long abundance associated with poorer overall survival in the TCGA-AML cohort. Our findings highlight a connection between SF3B1-dependent splicing, cell cycle progression, and tumorigenesis, offering new insights into the molecular mechanisms underlying cancer-associated splicing dysregulation.

## Background

Pre-mRNA splicing is facilitated by the spliceosome, a dynamic ribonucleoprotein complex composed of U1, U2, U4, U5, and U6 small nuclear ribonucleoproteins (snRNPs) along with numerous accessory proteins. Splicing occurs in a stepwise manner, beginning with the recognition of key sequence elements in the pre-mRNA, including the 5’ splice site, 3’ splice site, and branch point motif. The spliceosome undergoes conformational changes and catalytic activation to excise the intron and ligate the exons together, generating mature mRNA. SF3B1, a U2 snRNP-associated protein, plays a critical role in this process by stabilizing U2 binding to the branch point motif, ensuring proper recognition of the 3’ splice site. During spliceosome activation, SF3B1 undergoes phosphorylation at T313 and T434 by DYRK1A, which may influence its interaction with other spliceosomal components [1–4]. Additional phosphorylation sites at T244 and T248 have been identified through both low- and high-throughput approaches and are thought to promote interaction with the protein phosphatase PPP1R8 (NIPP1), suggesting a potential regulatory mechanism governing SF3B1 function [5, 6].

Recent studies have connected SF3B1 phosphorylation to cell cycle progression, with high phosphorylation levels observed in G2/M due to CDK2 activity, while lower phosphorylation levels are seen in G1/S due to the action of the PP2A and PP1 phosphatases [7], with specific phosphorylation primarily mediated by CDK1 [7, 8]. Additionally, PP1-mediated dephosphorylation of SF3B1 at T142p enables mitotic exit.

The critical role of SF3B1 in proper cell cycle progression is underscored by its knock-down, which leads to G2/M arrest [9–11]. Furthermore, SF3B1 inhibitors, also known as splicing inhibitors, induce cell cycle arrest at S and M phases [12, 13]. Notably, SF3B1 is the most commonly mutated component of the spliceosome in cancer. While the precise role of SF3B1 in cancer is still under investigation, its high prevalence in cancer cases strongly suggests its involvement in cell cycle progression [14, 15].

Non-driver mutations in SF3B1 are frequently observed in myelodysplastic syndromes (MDS) and other neoplasms affecting myeloid and lymphoid cells [16]. The most prevalent mutation is a heterozygous missense substitution known as K700E (SF3B1^K700E^). Additionally, SF3B1 mutations have been identified in various solid tumors, including uveal melanoma, breast cancer, pancreatic cancer, and prostate cancer [17]. This suggests that SF3B1 mutations promote cell cycle progression, in contrast to its knockdown or inhibition. Consequently, SF3B1 inhibitors have been proposed as potential therapeutic agents for cancer [18], with compounds like H3B-8800 and E7107 exhibiting a pronounced effect on SF3B1^K700E^ cells [19–21].

Phosphorylation of splicing factors has been shown to exert regulatory effects on protein-protein interactions both within the spliceosome and between splicing factors and pre-mRNA molecules [18]. Furthermore, it is well-established that alternative splicing undergoes dynamic regulation across the cell cycle phases [17, 22]. However, whether oncogenic SF3B1 reshapes splicing of cell-cycle regulators—and whether signalling pathways modulate those same exons—remains unclear.

Here, we delineate how the recurrent SF3B1^K700E^ mutation reshapes the splicing landscape of cell-cycle regulators in K562 cells, validate the robustness of key exon changes in an independent cellular context, and assess whether kinase–phosphatase signalling can modulate those same splice events. Our findings show that phosphorylation during the G2/M phase associated with splicing of ARPP19-long isoform and that ARPP19-long correlates with poor survival in acute myeloid leukemia (AML) patients and accelerates mitotic exit, suggesting a potential oncogenic role of this isoform. By integrating transcriptome profiling, targeted perturbations, and functional read-outs of the cell cycle, we uncover a spliceosomal programme that links SF3B1 mutation to G2/M control and to adverse outcome in acute myeloid leukaemia

## Material and methods

### Cell lines

HeLa (ATCC number: CRM-CCL-2) cells were grown in Dulbecco’s modified Eagle’s medium (DMEM). K562 (ATCC number: CCL-243TM), K562-WT and K562-SF3B1K700E cells were grown in Roswell Park Memorial Institute’s medium (RPMI-1640). Media was supplemented with 10% fetal bovine serum (FBS), 2 mM l-glutamine, 0.1 mg/ml penicillin and 0.1 mg/ml streptomycin. Cell lines were maintained at 37°C and 5% CO2 atmosphere.

### RNA sequencing and analysis

Total RNA samples for three biological replicates (approximately 1×10^6^ cells) from K562-WT and K562-SF3B1 K700E cells were subjected to sequencing. Raw reads were processed for quality trimming and adaptors removal using fastx_toolkit v0.0.14 and cutadapt v2.10. The processed reads were aligned to the human transcriptome and genome version GRCh37 using TopHat v2.1. Reads were counted with featureCounts version 2.0.1 using GENCODE v47. DESeq2 [23] was used to identify differentially expressed genes using FDR threshold of 0.1. rMATS (version 4.1.2) [24] was used to identify differential alternative splicing events. For each alternative splicing event, we used the calculation on both the reads mapped to the splice junctions and the reads mapped to the exon body [Junction Counts and Exon Counts (JCEC)] and filtered events by FDR threshold of 0.1. Gene set enrichment analysis was performed using the GeneAnalytics [25] online tool. FDR was controlled by the Benjamini-Hochberg procedure.

### RNA interference

A pool of four esiRNA oligomers against SF3B1 was purchased from Sigma-Aldrich (EHU131531). HeLa cells were grown to 20–30% confluence and transfected with 20 nM siRNA (esiSF3B1 or esiGFP for control) using TransIT-X2 transfection reagent (Mirus) following the manufacturer’s instructions. After 24 h of incubation, cell culture media was replaced and then incubated for additional 48-72 h. Knockdown efficiencies were verified by qPCR.

### qRT-PCR

RNA was isolated from cells using the GENEzol TriRNA Pure Kit (GeneAid). cDNA synthesis was carried out with the qScript cDNA Synthesis Kit (QuantaBio). qPCR was performed with the iTaq Universal SYBR Green Supermix (BioRad) on the Biorad iCycler. The comparative Ct method was employed to quantify transcripts, and delta Ct was measured in triplicate. Primers used in this study are provided in Supplementary Table S1.

### Cell cycle arrest

For G1/S synchronization, HeLa and K562 cells were cultured in DMEM with thymidine (T1895, Sigma-Aldrich) at a final concentration of 2 mM for 18 h. Cells were then washed with DMEM twice and cultured for 9 h in DMEM alone. Thymidine was then added at the same concentration and the cells were cultured for a further 18 h. For G2/M synchronization, HeLa and K562cells were cultured in DMEM with 40 ng/ml of nocodazole (NCO) (Sigma-Aldrich, M1404). Pharmacological inhibitors were added to G1/S or to G2/M synchronized cells as follows: okadaic acid (protein phosphatases inhibitor, Sigma-Aldrich, 39302), 20 nM for 12 h; INDY (DRKY1A inhibitor, Sigma-Aldrich, SML1011) 5 μM for 6 h. The cell cycle status was checked using flow cytometry.

### Immunoblotting

For immunoblotting, cells were harvested and lysed with RIPA lysis buffer supplemented with protease and phosphatase inhibitors, and 20 μg/μl of the extracts were run on a 10% TGX Stain-free FastCast gel (BioRad) and transferred onto a nitrocellulose membrane.

### Survival analysis

OncoSplicing [26] online analysis pipeline was used for splicing-based survival analysis and Kaplan-Mayer plot creation. Analysis was performed using SpliceSeq database PSI values and default optimal cutoffs. GEPIA [https://doi.org/10.1093/nar/gkx247] was used for expression-based survival analysis.

### Viral infection

ARPP19 and ENSA short and long isoforms were cloned into the p-Lenti-CMV-GFP vector (Addgene, plasmid # 17448). Lentiviral or retroviral particles were produced by transfecting HEK293T packaging cells with plasmids expressing ARPP19 and ENSA short or long isoforms and transfections were carried out using TransIT-X2 transfection reagent. HeLa cells were subsequently infected with the viral particles, and stable integrations were selected.

### Flow cytometry

Cells were trypsinized, washed with PBS, fixed overnight at −20°C with 70% ethanol in PBS, washed with PBS, and left for 30 min at 4°C. The cell suspension was then incubated with PBS containing 5 µg/mL DNase-free RNase and stained with propidium iodide (PI). Data were acquired using the LSRII Fortessa Analyzer machine at 10,000 events/sample. The percentage of cells in each cell-cycle phase was determined using Flow Jo software.

### Pharmacological inhibition of SF3B1

K562 cells were maintained in DMEM supplemented with 10% FBS and 1% penicillin-streptomycin at 37°C in a humidified 5% CO₂ incubator. Cells were seeded at a density of 1– 2 × 10⁵ cells/mL in culture plates and allowed to grow overnight. Pladienolide B was dissolved in dimethyl sulfoxide (DMSO) to prepare a 10 mM stock solution. For treatment, pladienolide B was diluted in culture medium to the desired working concentration (100 nM). Cells were treated with pladienolide B or an equivalent volume of DMSO (vehicle control) for 6 h.

## Results

### SF3B1 regulates splicing of cell cycle genes

SF3B1 function is critical for proper cell cycle progression. To search for splicing targets of SF3B1 with a pivotal role during the cell cycle, we conducted RNA-seq from WT and SF3B1^K700E^ K562 cells (n=3 independent cultures per genotype). Our analysis unveiled 22,136 alternative splicing events across 6,738 genes. These events encompassed various types, with 40% involving exon exclusion (8,953 events), 5% alternative 5’ splice sites (1,098 events), 16% alternative 3’ splice sites (3,583 events), 33% mutually exclusive exons (7,408 events), and 4% intron retention (1,094 events) (Fig. 1A and S1A, FDR < 5%, rMATS, Supplementary Table S2). Our findings are consistent with prior reports of cryptic 3′ splice-site usage in SF3B1^K700E^-induced chronic lymphocytic leukemia (CLL) and K562 cells [27].

**Figure 1.**
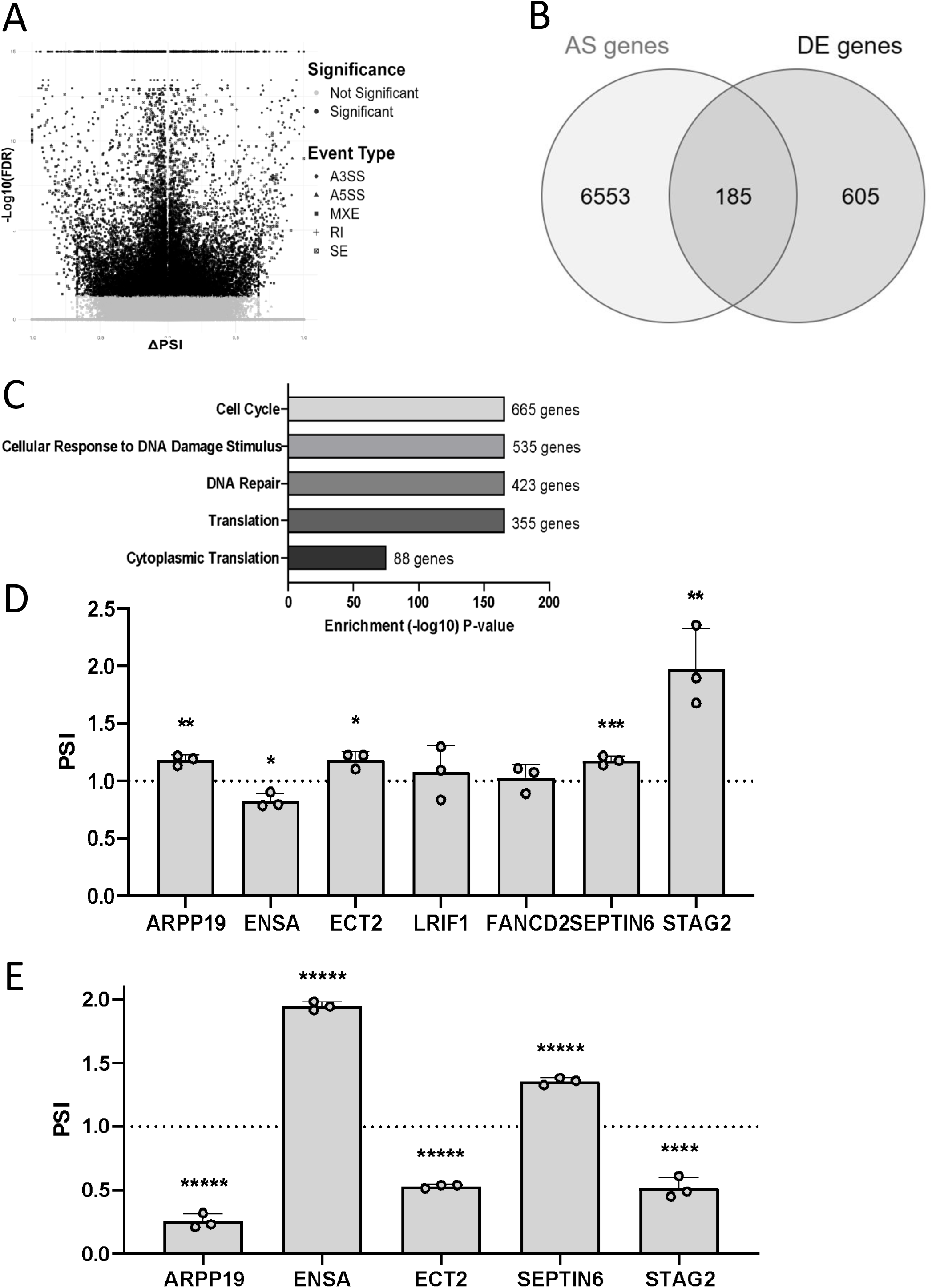
SF3B1 regulates alternative splicing of cell cycle checkpoint genes. **A-C.** rMATs analysis was done on RNA-seq data from WT K562 or SF3B1^K700E^. A volcano plot showing differential splicing events between WT K562 cells and K562^K700E^. The x-axis represents the inclusion level difference (ΔPSI), and the y-axis shows statistical significance (-log10 p-value). Significant events (FDR < 0.05) are in black, while non-significant ones are in gray **(A)**. Venn diagram representing the overlap between differentially expressed genes (DE genes) and alternatively spliced genes (AS genes) **(B)**. Gene set enrichment analysis was performed on alternatively spliced genes comparing WT K562 cells and K562^K700E^ cells using the GeneAnalytics tool [26] (**C)**. **D.** RNA was extracted from WT K562 and K562^K700E^ cells, converted to cDNA and RT-PCR was performed. RT-PCR analysis of the PSI (percentage splice in) relative to RT-PCR analysis of total mRNA for ARPP19 exon 2 inclusion, ENSA exon 3 inclusion, ECT2 exon 5 inclusion, LRIF1 exon 2 inclusion, FANCD2 exon 22 inclusion, SEPTIN6 exon 10 inclusion and STAG2 exon 31 inclusion. **E**. Cells were transfected with siGFP or siSF3B1 for 72 h. RNA was extracted, converted to cDNA and RT-PCR was performed. RT-PCR analysis of the PSI relative to RT-PCR analysis of total mRNA for ARPP19 exon 2 inclusion, ENSA exon 3 inclusion, ECT2 exon 5 inclusion, LRIF1 exon 2 inclusion, FANCD2 exon 22 inclusion, SEPTIN6 exon 10 inclusion and STAG2 exon 31 inclusion. Values represent averages of three independent experiments ±SD *p<0.05; **p<0.01; ***p<0.005; ****p<0.001; Student’s t-test, compared to PSI and total mRNA values for cells transfected with siGFP (y=1).

Gene expression analysis identified 790 differentially expressed genes, with 599 upregulated genes in SF3B1^K700E^ and 191 downregulated genes. Upregulated genes were enriched for Akt signaling (34 genes), angiogenesis (11 genes) and other signaling pathways while downregulated genes showed enrichment for immune and metabolic pathways (Fig. S1B). 185 differentially expressed genes are also alternatively spliced (P < 0.00001, Fisher’s exact test, Fig. 1B, Supplementary Table S3).

From the 6,738 genes regulated in alternative splicing, we found enrichment for genes involved in cell cycle (665 genes), DNA damage response (535 genes) and translation (355 genes) (Fig. 1C). To elucidate the role of SF3B1 during the cell cycle, we selected seven alternative splicing events in cell cycle checkpoint genes for validation (Fig. S1C-H). Out of the seven events chosen, we validated five (Fig. 1D and S2A); STAG2, ENSA, and ARPP19, which play roles in the M-phase, and SEPTIN6 and ECT2, which are involved in the S-phase.

To further confirm that SF3B1 regulates the splicing of the five previously validated events, we silenced SF3B1 in three biological replicates of HeLa cervical cancer cells using siRNA (Supplementary Fig. S2B). Given that SF3B1^K700E^ is a gain-of-function mutation, we hypothesized that silencing SF3B1 would produce an opposite effect on splicing. Indeed, we observed the opposite effect for STAG2, ENSA, ARPP19, and ECT2. SEPTIN6 exhibited mild exon 10 inclusion in SF3B1^K700E^ K562 cells, and a similar trend was seen in HeLa cells upon SF3B1 silencing (20% reduction, Fig. 1E). Additionally, we observed a decrease in total mRNA levels for ENSA, ARPP19, ECT2, and SEPTIN6 in HeLa cells, which was not seen in SF3B1^K700E^ (Supplementary Fig. S2C). These findings further support the role of SF3B1 in regulating alternative splicing of key cell cycle genes and note a potential alteration of total mRNA levels by SF3B1.

### Kinase and phosphatase signaling modulate SF3B1-linked splice events

To test whether signaling pathways alter SF3B1^K700E^ splicing targets, we synchronized K562 and HeLa cells in the G1/S and G2/M phases and analyzed the splicing events regulated by SF3B1 (Fig. 2A-H). Results were consistent in both cell lines for ARPP19 exon 2 inclusion, which gives rise to ARPP19-long isoform, with respectively 60% and 40% reduction in G1/S in K562 and Hela cells (Fig. 2A&E). In G2/M, we saw a 40% increase in ARPP19-long in both cell lines (Fig. 2C&G). Total mRNA amount of ARPP19 and ENSA decreased in K562 G2/M cells (Fig. S3B). These findings indicate that inclusion of ARPP19 exon 2 varies with cell-cycle phase.

**Figure 2.**
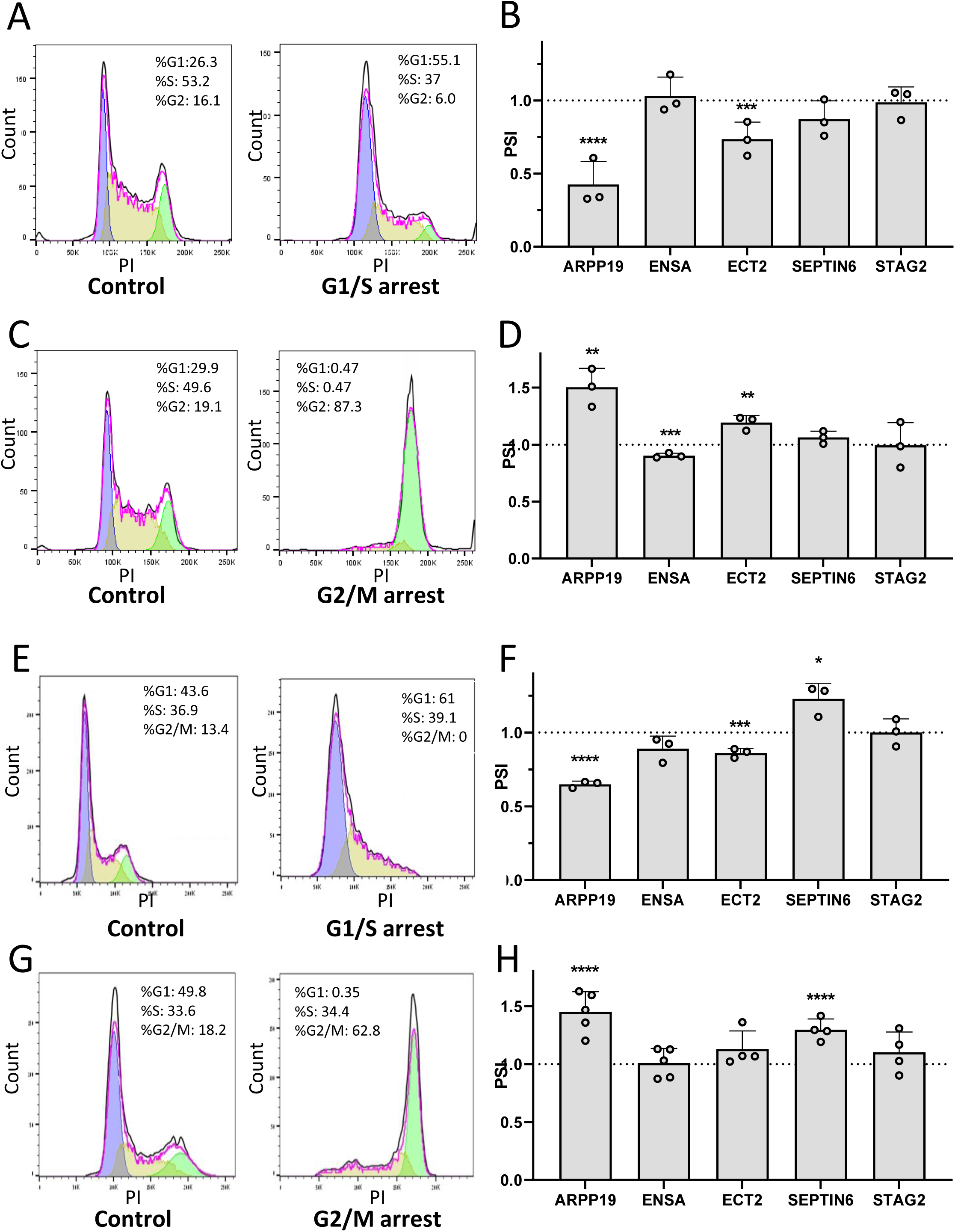
Effect of cell cycle stage on alternative splicing. **A-B**. K562 cells were treated with thymidine at a final concentration of 2 mM, for 18 h. Cells were washed then after 9 h treated again with 2 mM thymidine. After 18 h cells were collected and analysed by flow cytometry (**A**) and RT-PCR (**B**). RNA was extracted, converted to cDNA and RT-PCR analysis of the PSI relative to RT-PCR analysis of total mRNA for ARPP19 exon 2 inclusion, ENSA exon 3 inclusion, ECT2 exon 5 inclusion, SEPTIN6 exon 10 inclusion and STAG2 exon 31 inclusion was performed. Values represent averages of three independent experiments ±SD *p<0.05; ***p<0.005; ****p<0.001; Student’s t-test, compared to PSI for control cells. **C-D**. K562 cells were treated with 40 ng/ml nocodazole (NCO) for 20 h. Cells were collected and analysed by flow cytometry (**C**) and RT-PCR (**D**). RNA was extracted, converted to cDNA and RT-PCR analysis of the PSI relative to RT-PCR analysis of total mRNA for ARPP19 exon 2 inclusion, ENSA exon 3 inclusion, ECT2 exon 5 inclusion, SEPTIN6 exon 10 inclusion and STAG2 exon 31 inclusion was performed. Values represent averages of three independent experiments ±SD ****p<0.001; Student’s t-test, compared to PSI for control cells. **E-F**. HeLa cells were treated with thymidine at a final concentration of 2 mM, for 18 h. Cells were washed then after 9 h treated again with 2 mM thymidine. After 18 h cells were collected and analysed by flow cytometry (**E**) and RT-PCR (**F**). RNA was extracted, converted to cDNA and RT-PCR analysis of the PSI relative to RT-PCR analysis of total mRNA for ARPP19 exon 2 inclusion, ENSA exon 3 inclusion, ECT2 exon 5 inclusion, SEPTIN6 exon 10 inclusion and STAG2 exon 31 inclusion was performed. Values represent averages of three independent experiments ±SD *p<0.05; ***p<0.005; ****p<0.001; Student’s t-test, compared to PSI for control cells. **G-H**. HeLa cells were treated with 40 ng/ml nocodazole (NCO) for 20 h cells were collected and analysed by flow cytometry (**G**) and RT-PCR (**H**). RNA was extracted, converted to cDNA and RT-PCR analysis of the PSI relative to RT-PCR analysis of total mRNA for ARPP19 exon 2 inclusion, ENSA exon 3 inclusion, ECT2 exon 5 inclusion, SEPTIN6 exon 10 inclusion and STAG2 exon 31 inclusion was performed. Values represent averages of three independent experiments ±SD ****p<0.001; Student’s t-test, compared to PSI for control cells.

We next asked whether kinase signaling can modulate the splice events linked to SF3B1^K700E^. This analysis encompassed both synchronized and unsynchronized cells to discern the significance of phosphorylation at specific cell cycle stages. We started with the kinase DYRK1A, known to have a role in G1/S and to phosphorylate SF3B1 on T434 [4]. To this end, we arrested K562 and HeLa cells in G1/S and upon release, cells were treated with the DYRK1A/B inhibitor INDY for 6 h. FACS analysis revealed that the treatment did not affect cell cycle progression in naïve cells or following arrest (Fig. 3A&D), a critical observation considering the sensitivity of splicing of certain targets to the cell cycle stage. Our results show that DYRK1A inhibition in naïve cells or in G1/S reduces the amount of ARPP19-long by around 30% in both cell lines (Fig. 3B&E and S4A-P). In addition, we observed a mild reduction in ENSA exon 3 inclusion in HeLa cells (Fig. 3F). This result is consistent with DYRK1A activity modulating ARPP19 exon 2 inclusion.

**Figure 3.**
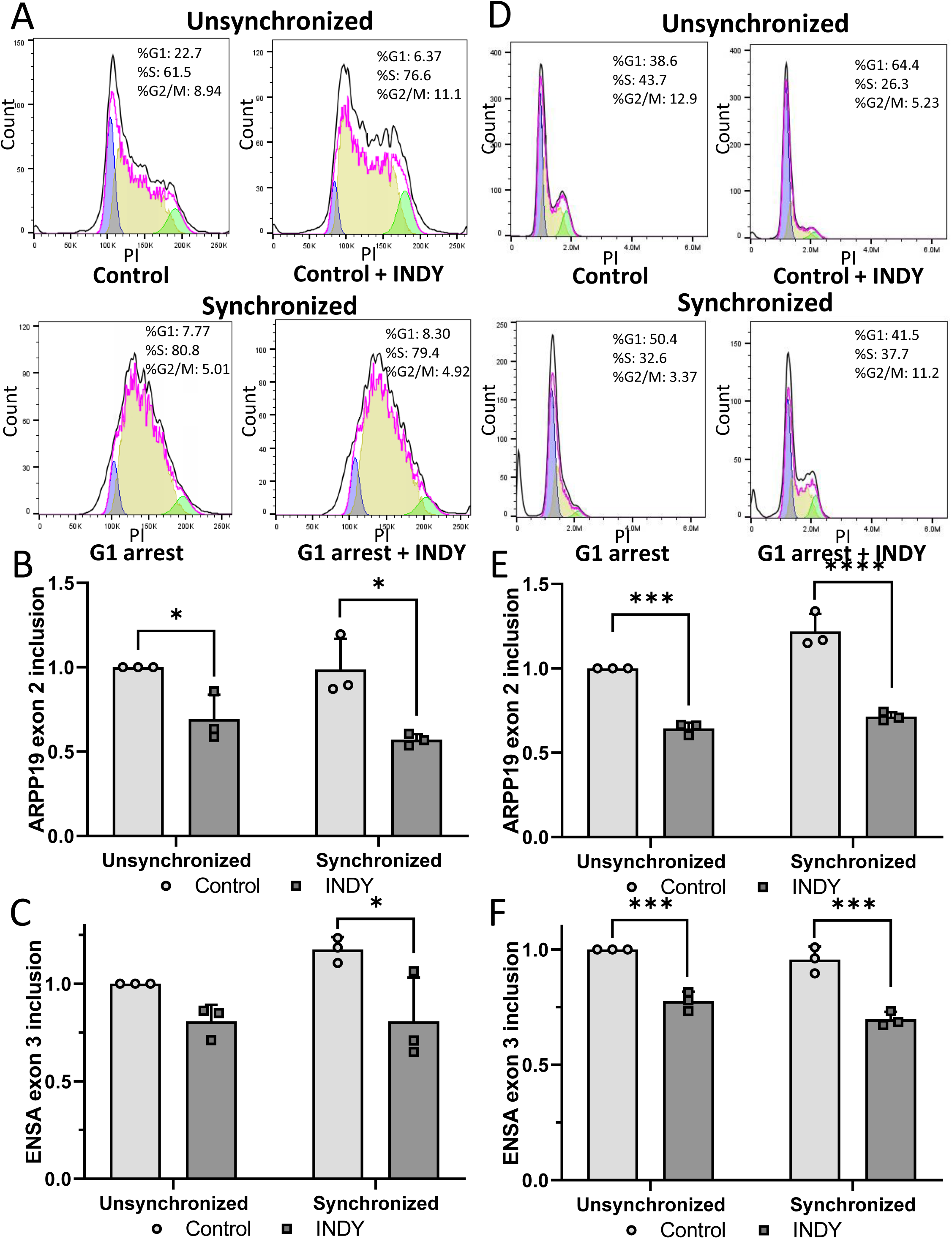
The effect of inhibition of DYRK1A, following G_1_/S synchronization, on alternative splicing. A-C. K562 cells were treated with thymidine at a final concentration of 2 mM, for 18 h. Cells were washed then after 9 h treated again with 2 mM thymidine. After 18 h cells were washed and then treated with media or the DYRK1A inhibitor INDY at a final concentration of 5 μM for 6 h. Cells were collected for analysis by flow cytometry, using propidium iodide (PI) to measure DNA content. The proportions of cells accumulated in S-phase is shown as a fold-change relative to untreated cells and the results presented as histograms of the various cell treatment groups: non-treated (control) cells (**A**), thymidine-treated cells (**B**), thymidine treated cells, after 6 h further incubation with DMEM alone (**C**), thymidine-treated cells, after 6 h incubation with INDY (**D**). **E-F**. Cells were collected for RT-PCR analysis. RNA was extracted, converted to cDNA and RT-PCR analysis of the PSI relative to RT-PCR analysis of total mRNA of ARPP19 exon 2 inclusion (**E**), ENSA exon 3 inclusion (**F).** Values represent averages of three independent experiments ±SD *p<0.05; Student’s t-test. **G-J.** HeLa cells were treated with thymidine at a final concentration of 2 mM, for 18 h. Cells were washed then after 9 h treated again with 2 mM thymidine. After 18 h cells were washed and then treated with media or the DYRK1A inhibitor INDY at a final concentration of 5 μM for 6 h. Cells were collected for analysis by flow cytometry, using propidium iodide (PI) to measure DNA content. The proportions of cells accumulated in S-phase is shown as a fold-change relative to untreated cells and the results presented as histograms of the various cell treatment groups: non-treated (control) cells (**G**), thymidine-treated cells (**H**), thymidine treated cells, after 6 h further incubation with DMEM alone (**I**), thymidine-treated cells, after 6 h incubation with INDY (**J**). **K-L**. Cells were collected for RT-PCR analysis. RNA was extracted, converted to cDNA and RT-PCR analysis of the PSI relative to RT-PCR analysis of total mRNA of ARPP19 exon 2 inclusion (**K**), ENSA exon 3 inclusion (**L).** Values represent averages of three independent experiments ±SD *p<0.05; ***p<0.005; ****p<0.001; Student’s t-test.

Protein-phosphatase 2A (PP2A) counteracts several mitotic kinases and has been reported to de-phosphorylate SF3B1 during the M-to-G1 transition. To investigate its effect on splicing, we used its inhibitor, okadaic acid, in both naïve K562 and HeLa cells, as well as following synchronization at G1/S. We began by arresting the cells in G1 phase and, upon release, treated them with okadaic acid for 12 h. Our results show a 70% increase in ARPP19 exon 2 inclusion in unsynchronized cells (predominantly in G1) and a 20% increase in ARPP19 exon 2 inclusion in synchronized cells (enriched for G2/M) (Fig. 4A-E and S5A-Q). Although we measured splicing changes in five genes, our results were particularly robust for ARPP19. Taken together, the DYRK1A- and PP2A-inhibitor data show that kinase– phosphatase signaling can shift the inclusion of ARPP19 exon 2 and other SF3B1-responsive exons, reinforcing a model in which a phosphorylation-sensitive spliceosome checkpoint fine-tunes cell-cycle gene splicing.

**Figure 4.**
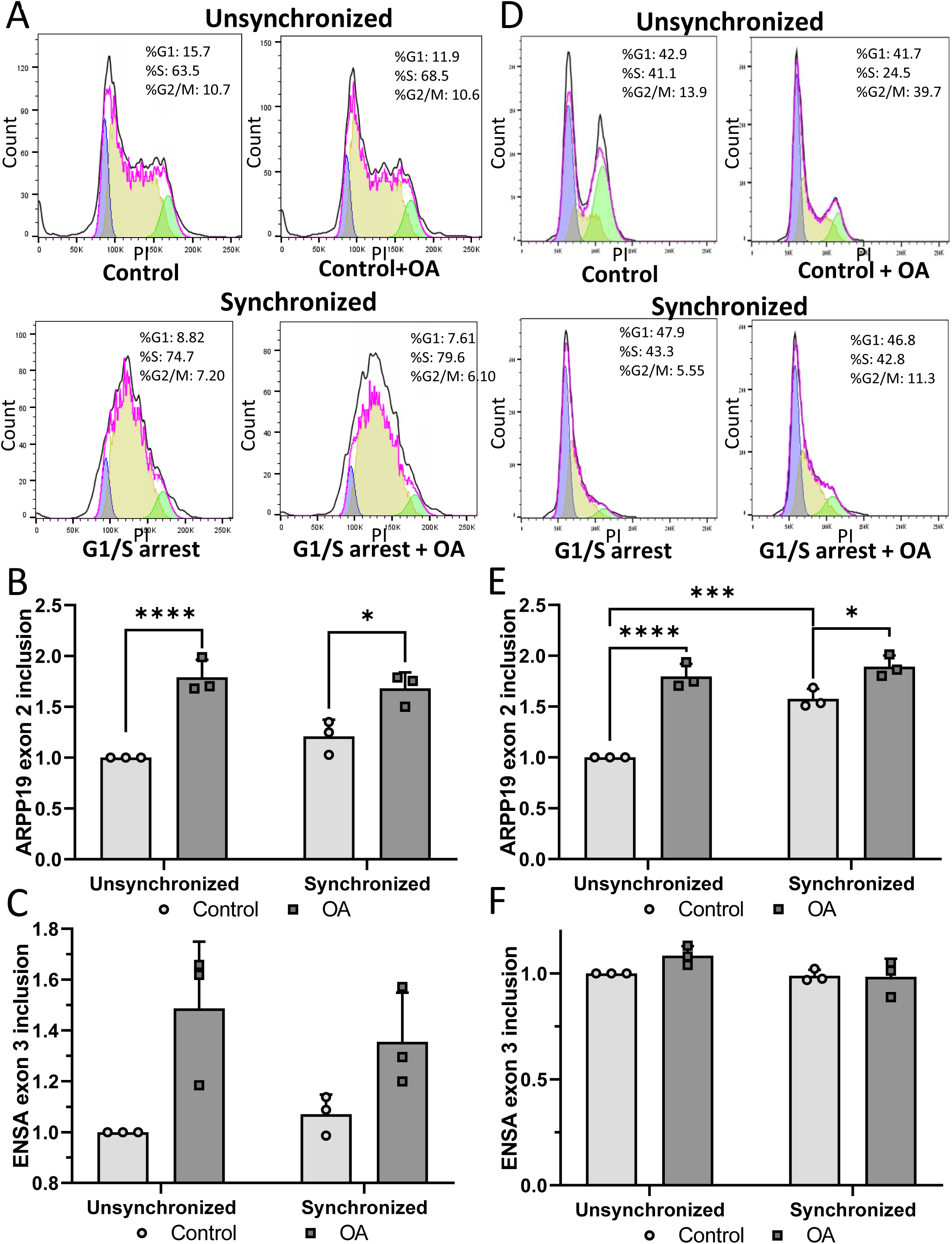
The effect of inhibition of PP2A, following G_1_/S synchronization, on alternative splicing. A-F. K562 cells were treated with thymidine at a final concentration of 2 mM, for 18 h. Cells were washed then after 9 h treated again with 2 mM thymidine. After 18 h cells were washed and then treated with media or the PP2A inhibitor okadaic acid (OA) at a final concentration of 20 nM for 12 h. **A-D.** Cells were collected for analysis by flow cytometry using PI to measure DNA content. The amount of cells accumulated in S-phase is shown as a fold-change relative to untreated cells and the results presented as histograms of the various cell treatment groups: non-treated (control) cells (**A**), OA treatment for 12 h (**B**) thymidine-treated cells (**C**), thymidine-treated cells, plus OA treatment for 12 h (**D**). **E-F**. Cells were collected for RT-PCR. RNA was extracted, converted to cDNA and RT-PCR analysis of the PSI relative to RT-PCR analysis of total mRNA of ARPP19 exon 2 inclusion (**E**), ENSA exon 3 inclusion (**F**). Values represent averages of three independent experiments ±SD *p<0.05; ****p<0.001; Student’s t-test. **G-M**. HeLa cells were treated with thymidine at a final concentration of 2 mM, for 18 h. Cells were washed then after 9 h treated again with 2 mM thymidine. After 18 h cells were washed and then treated with media or the PP2A inhibitor okadaic acid (OA) at a final concentration of 20 nM for 12 h. **G-K.** Cells were collected for analysis by flow cytometry using PI to measure DNA content. The amount of cells accumulated in S-phase is shown as a fold-change relative to untreated cells and the results presented as histograms of the various cell treatment groups: non-treated (control) cells (**G**), OA treatment for 12 h (**H**) thymidine-treated cells (**I**), thymidine-treated cells, plus 12 h with DMEM (**J**), thymidine-treated cells, plus OA treatment for 12 h (**K**). **L-M**. Cells were collected for RT-PCR. RNA was extracted, converted to cDNA and RT-PCR analysis of the PSI relative to RT-PCR analysis of total mRNA of ARPP19 exon 2 inclusion (**L**), ENSA exon 3 inclusion (**M**). Values represent averages of three independent experiments ±SD *p<0.05; ****p<0.001; Student’s t-test.

### Alternative splicing of ARPP19 is associated with AML patient survival

We hypothesize that SF3B1^K700E^ is associated with alternative splicing of cell-cycle regulators that may influence disease course. To evaluate whether the isoforms linked to SF3B1 could serve as a prognostic factor for AML, we performed a survival analysis of SF3B1 splicing targets using clinical RNA-seq datasets from TCGA [28, 29]. Our results indicate that ARPP19 isoform-based survival analysis serves as a significant predictor for prognosis while gene expression-based analysis does not (Fig. 5A&B and S6A-F). In contrast, low expression of ENSA serves as a survival predictor, while its exon 3 splicing does not (Fig. 5C&D). Specifically, in AML, increased inclusion of ARPP19 exon 2, resulting in the long isoform (ARPP19-long), correlates with poorer survival compared to ARPP19-short. We observed that ARPP19-long increases in G2/M and tracks with splice changes seen under kinase-dominated conditions. Next, we aimed to investigate the role of this isoform in the cell cycle.

**Figure 5.**
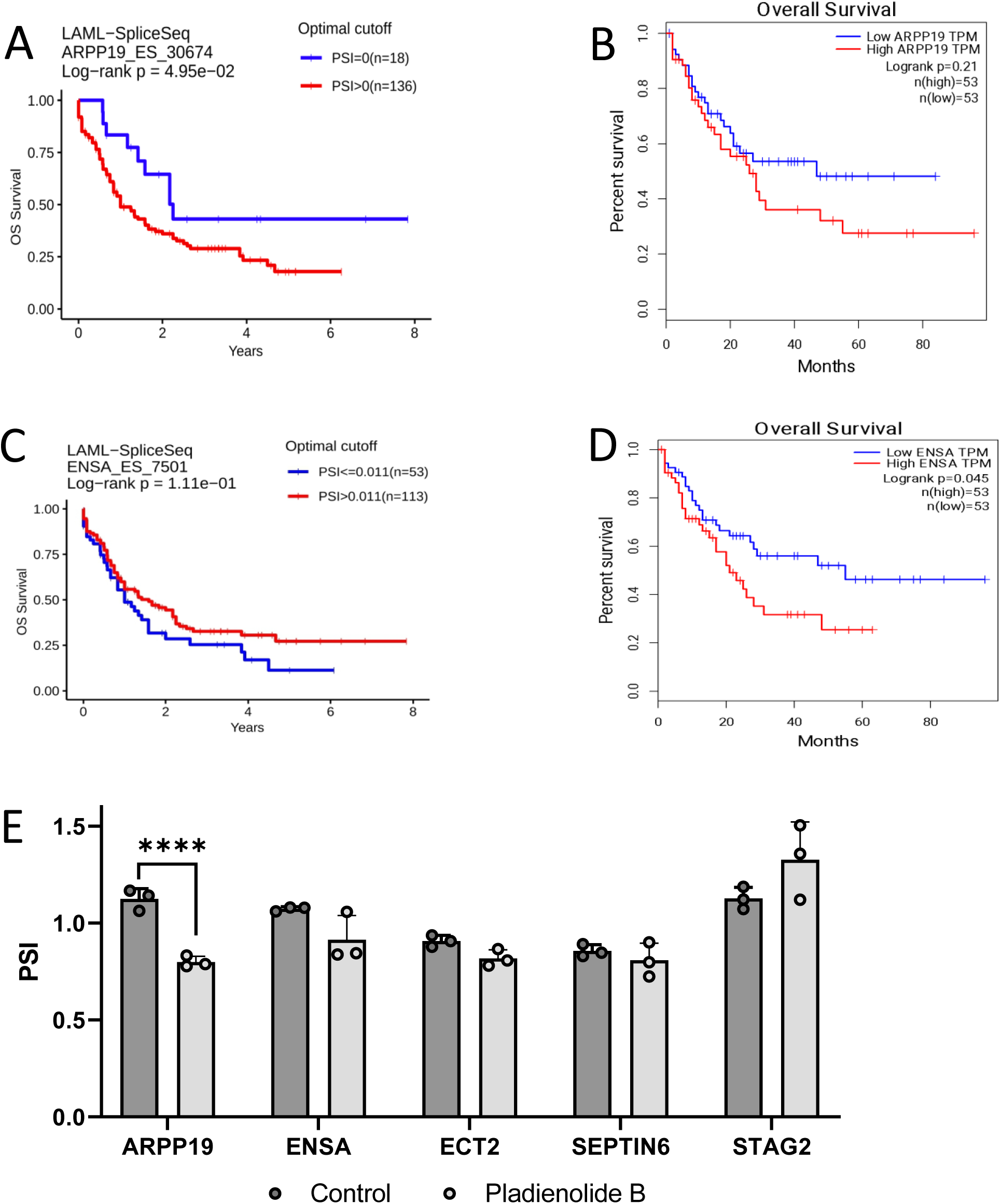
Survival analysis of the SF3B1 splicing targets using clinical RNA-seq datasets from TCGA for AML patients. **A-B.** Kaplan-Meier curves for total mRNA of ENSA (**A**); and for ENSA exon 3 inclusion (**B**). High exon inclusion level is colored in red and lower inclusion level in blue. Error bars represent 95% confidence intervals of the mean. **C**-**D**. Kaplan-Meier curves for total mRNA of ARPP19 (**C**); and for ARPP19 exon 2 inclusion (**D**). High exon inclusion level is colored in red and lower inclusion level in blue. Error bars represent 95% confidence intervals of the mean. **E.** K562 cells were treated with pladienolide B, an SF3B1 inhibitor, at 100 nM for 24 h. Cells were collected for RT-PCR. RNA was extracted, converted to cDNA and RT-PCR analysis of the PSI relative to RT-PCR analysis of total mRNA of ENSA exon 3 inclusion, ARPP19 exon 2 inclusion, ECT2 exon 5 inclusion, SEPTIN6 exon 10 inclusion and STAG2 exon 31 inclusion. Values represent averages of three independent experiments ±SD *p<0.05; ****p<0.001; Student’s t-test.

### SF3B1 inhibition by pladienolide B reduces the ARPP19-long isoform

Given the correlation between ARPP19-long isoform and poor AML survival, we next examined whether SF3B1 inhibition could influence ARPP19 splicing, as it is speculated to increase survival [13]. Since SF3B1^K700E^ promotes ARPP19 exon 2 inclusion (Fig. 1C), we hypothesized that pharmacological inhibition of SF3B1 would have the opposite effect, leading to a reduction in ARPP19-long isoform expression, similar to SF3B1 knockdown (Fig. 1E). To test this, we treated K562 cells with the well-characterized SF3B1 inhibitor pladienolide B (100 nM) for 6 h, and assessed splicing of STAG2, ENSA, ARPP19, ECT2, and SEPTIN6. Our results show a decrease in ARPP19 exon 2 inclusion (Fig. 5E), meaning less ARPP19-long, highlighting a tractable splice event whose modulation may complement existing therapeutic strategies in AML.

### ARPP19-long is associated with faster mitotic exit

Our findings underscore ARPP19 as a target of SF3B1, with its isoforms associated with AML survival. To gain further insights, we sought to elucidate the distinct functions of ARPP19 isoforms. ARPP19 forms a transient inhibitory complex with PP2A-B55, with its heterodimeric partner, ENSA, also identified as a target of SF3B1. Interestingly, however, the isoforms of ENSA was not significantly associated with survival in the same cohort. ARPP19 and ENSA act cooperatively to restrain PP2A-B55 during mitotic entry. Although Cyclin B/CDK1 activation initiates mitosis, its full effect depends on the inhibition of PP2A-B55 activity [30]. Taken together, these observations underscore the potential regulatory contribution of ARPP19 and ENSA to precise timing of mitotic progression.

To understand the differential function of ARPP19 and ENSA isoforms at this stage, we expressed each of the protein isoforms in HeLa cells. We generated stable HeLa cells with GFP-empty, GFP-ARPP19-long, or GFP-ARPP19-short (Fig. 6A), as well as GFP-empty, GFP-ENSA-long, or GFP-ENSA-short (Fig. S7A). We then synchronized the cells in G2/M following 20 h with nocodazole (Fig 7B and S7B). After releasing the cells, we monitored their cell cycle stage after 3 h. Our results demonstrate a differential role for both ARPP19 and ENSA isoforms during mitotic exit (Fig. 6B&C and S7B&C). Specifically, cells overexpressing ARPP19-long progressed to G1 faster than those overexpressing ARPP19-short, while cells overexpressing ENSA-short progressed to G1 more slowly than those overexpressing ENSA-long (Fig. 6C and Supplementary Fig. S7C). These distinct behaviors are consistent with the splice shifts driven by SF3B1^K700E^ —namely increased ARPP19-long and ENSA-short—and with the clinical association between ARPP19-long abundance and poorer AML survival. Collectively, the data indicate that the SF3B1-linked isoform pattern can alter the timing of mitotic exit, offering a potential cellular pathway through which SF3B1 mutations contribute to oncogenic phenotypes.

**Figure 6.**
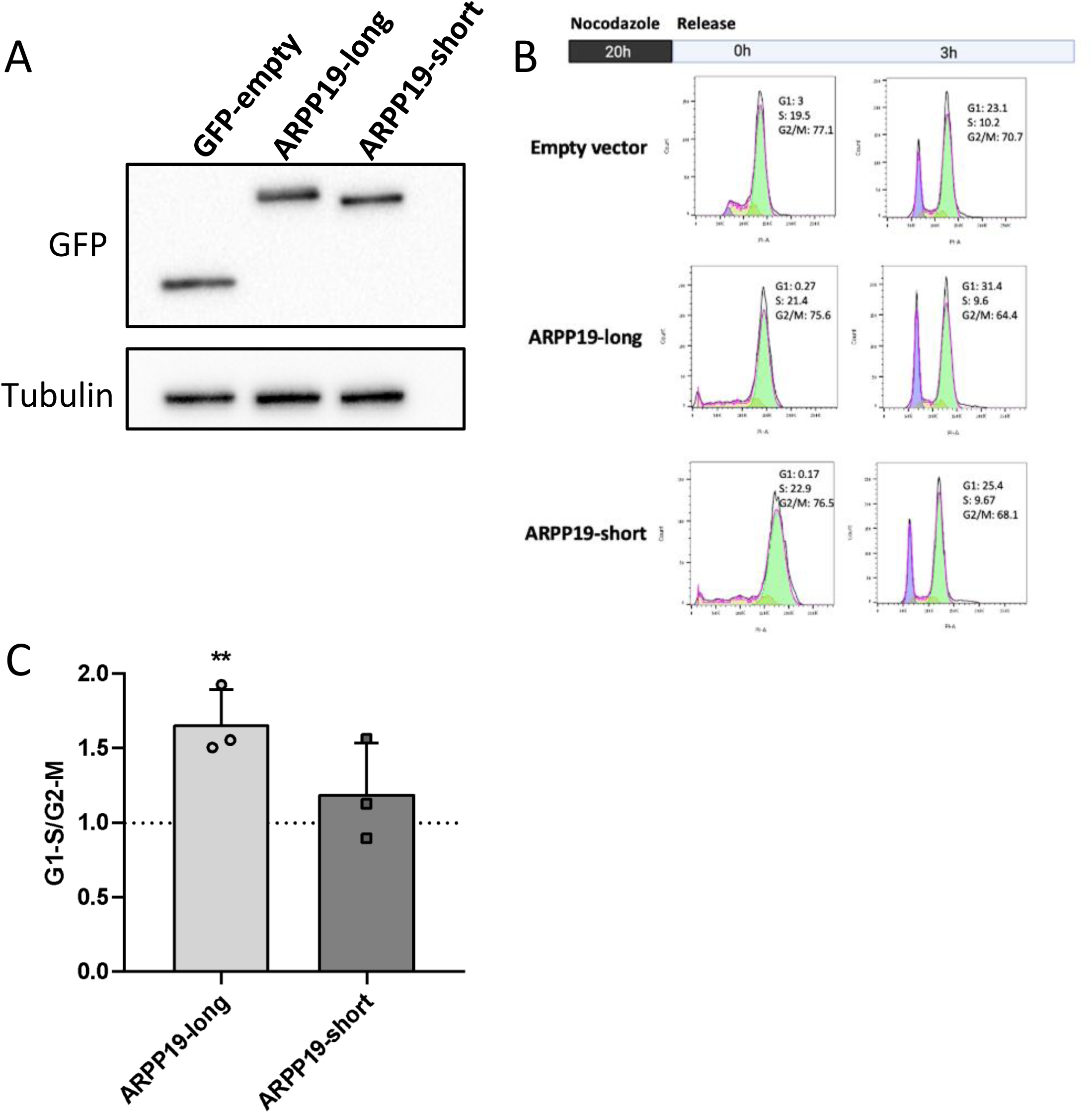
ARPP19 isoform function. ARPP19-short/long were cloned to pLenti CMV GFP vector and lentiviral particles were produced by expressing ARPP19 short or long isoform in the HEK293T packaging cell-line using TransIT-X2 transfection reagent. HeLa cells were infected with viral particles to overexpress either the short or long isoform of ARPP19. Protein was collected, and immunoblotting was conducted with the indicated antibodies and quantified relative to the housekeeping protein tubulin (**A**). The stable cell lines were arrested at G2/M using NCO for 20 h, then the cells were released for 3 h, collected and analyzed by flow cytometry using PI to measure DNA content (**B**), and the analysis was blotted in bar graph **(C).**

## Discussion

Given that nearly all human genes undergo splicing, this process is indispensable for maintaining cellular homeostasis. Among the many cellular processes dependent on splicing, the cell cycle stands out due to its strict requirement for timely and coordinated gene expression. Alterations in cell cycle regulation drive cancer, and SF3B1, a key spliceosome component, has been implicated in this process. SF3B1 mutations and phosphorylation changes have been linked to aberrant splicing, leading to disrupted cell cycle control and tumorigenesis. In this study, we show that the hotspot mutation SF3B1^K700E^ rewires a discrete set of splice events enriched for G2/M regulators; that inclusion of key splicing events, most notably ARPP19 exon 2, varies with cell-cycle phase; and that these same events respond to kinase and phosphatase perturbations, consistent with a phosphorylation-sensitive spliceosome checkpoint. By mapping these associations and probing their functional consequences in cultured cells, we outline how SF3B1-linked splicing changes—particularly the shift toward the ARPP19-long isoform—could influence mitotic timing and, by extension, cancer progression.

The functional relationship between ARPP19 and ENSA further supports the idea that SF3B1-mediated splicing contributes to mitotic regulation in a coordinated manner. Both genes likely evolved from a common ancestor, given their structural similarities and overlapping roles in PP2A-B55 inhibition. Interestingly, while ARPP19 alternative splicing correlates with AML patient survival, ENSA splicing does not, despite both genes functioning within the same PP2A-B55 regulatory complex. This suggests that ARPP19-long may have an additional moonlighting role in cancer progression, potentially acting as a key driver of mitotic exit dysregulation. In addition, our results suggest that phosphorylated SF3B1 and SF3B1^K700E^ promotes ARPP19-long and ENSA-short (Fig. 1D and 3B&E), both of which facilitate faster mitotic exit. This raises the possibility that cancer cells exploit SF3B1 mutations to shift the balance toward isoforms that accelerate cell cycle progression, ultimately supporting tumor growth. Importantly, our findings demonstrate that inhibition of SF3B1, such as treatment with pladienolide B, can reverse these splicing alterations, reducing the levels of ARPP19-long. This suggests that splicing modulation could serve as a therapeutic strategy, particularly for malignancies like AML, where SF3B1-driven splicing patterns correlate with poor prognosis.

## Data availability

The data generated in this study are currently in the process of being submitted to GEO. Once the submission is complete, access details will be provided.

## Acknowledgments

We thank Liran Shlush for K562-WT and K562-SF3B1^K700E^ cells. The results shown here are in part based upon data generated by the TCGA Research Network: https://www.cancer.gov/tcga.

## Author contribution

MB, EE, AS, MP: Conceptualization, validation, formal analysis, investigation, visualization, methodology, and writing-review-editing. SJH, OG: Validation. MSu, MB: Resources. GK: Visualization and project administration. MS: Conceptualization, funding acquisition, supervision, and writing-original draft.

## Supplementary Figures

**Sup. Figure 1: A.** rMATS analysis for the differential alternative splicing was performed on RNA-seq of WT K562 or SF3B1^K700E^. **B-G** RPKM for each alignment track are shown as sashimi plots. Arcs denote splice junctions, quantified in spanning reads, as specified near each exon junction of ARPP19 **(B)**, ECT2 **(C)**, LRIF1 **(D)**, FANCD2 **(E)**, STAG2 **(F)**, SEPTIN6 **(G)**. **H.** Schematic representation of primer design for detecting alternative splicing events for ARPP19, ENSA, ECT2, SEPTIN6 and STAG2 genes. Primers (black arrows) were positioned in flanking constitutive exons to amplify and distinguish between splice isoforms. Exons are shown as pink boxes, introns as black lines, and alternative splicing events.

**Sup. Figure 2. A.** RNA was extracted from WT K562 and K562^K700E^ cells, converted to cDNA and RT-PCR was performed. RT-PCR analysis of total mRNA of ARPP19, ENSA, ECT2, LRIF1, FANCD2, SEPTIN6 and STAG2 relative to CycloA and hTBP reference genes. Values represent averages of three independent experiments ±SD *p<0.05; **p<0.01; ***p<0.005; ****p<0.001; Student’s t-test. **B-C**. Cells were transfected with siGFP or siSF3B1 for 72 h. RNA was extracted, converted to cDNA and RT-PCR was performed. **(B)** ARPP19, ENSA, ECT2, LRIF1, FANCD2, SEPTIN6 and STAG2 relative to CycloA and hTBP reference genes **(C)**. Values represent averages of three independent experiments ±SD *p<0.05; **p<0.01; ***p<0.005; ****p<0.001; Student’s t-test, compared to total mRNA values for cells transfected with siGFP (y=1).

**Sup. Figure 3. A**. K562 cells were treated with thymidine at a final concentration of 2 mM, for 18 h. Cells were washed, then after 9 h treated again with 2 mM thymidine. After 18 h cells were collected and analysed by RT-PCR. RNA was extracted, converted to cDNA and RT-PCR analysis of the total mRNA relative to CycloA and hTBP reference genes for ARPP19, ENSA, ECT2, SEPTIN6 and STAG2 was performed. Values represent averages of three independent experiments ±SD *p<0.05; Student’s t-test, compared to total mRNA for control cells. **B**. K562 cells were treated with 40 ng/ml nocodazole (NCO) for 20 h cells were collected and analyzed by RT-PCR. RNA was extracted, converted to cDNA and RT-PCR analysis of the total mRNA relative to CycloA and hTBP reference genes for ARPP19, ENSA, ECT2, SEPTIN6 and STAG2 was performed. Values represent averages of three independent experiments ±SD *p<0.01; Student’s t-test, compared to total mRNA for control cells. **C.** HeLa cells were treated with thymidine at a final concentration of 2 mM, for 18 h. Cells were washed, then after 9 h treated again with 2 mM thymidine. After 18 h cells were collected and analyzed by RT-PCR. RNA was extracted, converted to cDNA and RT-PCR analysis of the total mRNA relative to CycloA and hTBP reference genes for ARPP19, ENSA, ECT2, SEPTIN6 and STAG2 was performed. Values represent averages of three independent experiments ±SD *p<0.05; Student’s t-test, compared to total mRNA for control cells. **D**. HeLa cells were treated with 40 ng/ml NCO for 20 h, then cells were collected and analyzed by RT-PCR. RNA was extracted, converted to cDNA and RT-PCR analysis of the total mRNA relative to CycloA and hTBP reference genes for ARPP19, ENSA, ECT2, SEPTIN6 and STAG2 was performed. Values represent averages of three independent experiments ±SD *p<0.01; Student’s t-test, compared to total mRNA for control cells.

**Sup. Figure 4. A-H.** K562 cells were treated with thymidine at a final concentration of 2 mM, for 18 h. Cells were washed, then after 9 h, treated again with 2 mM thymidine. After 18 h cells were washed and then treated with media or the DYRK1A inhibitor INDY at a final concentration of 5 μM for 6 h. Cells were collected for RT-PCR analysis. RNA was extracted, converted to cDNA and total mRNA relative to CycloA and hTBP reference genes is shown for ARPP19 (**A**), ENSA (**B**). RT-PCR analysis of the PSI relative to RT-PCR analysis of total mRNA of ECT2 exon 5 inclusion (**C&D**), SEPTIN6 exon 10 inclusion (**E&F**), and STAG2 exon 31 inclusion (**G&H**) was performed. Values represent averages of three independent experiments ±SD *p<0.05; Student’s t-test compared to PSI and total mRNA for control cells. **I-P.** HeLa cells were treated with thymidine at a final concentration of 2 mM, for 18 h. Cells were washed, then after 9 h, treated again with 2 mM thymidine. After 18 h cells were washed and then treated with media or the DYRK1A inhibitor INDY at a final concentration of 5 μM for 6 h. Cells were collected for RT-PCR analysis. RNA was extracted, converted to cDNA and total mRNA relative to CycloA and hTBP reference genes is shown for ARPP19 (**I**), ENSA (**J**). RT-PCR analysis of the PSI relative to RT-PCR analysis of total mRNA of ECT2 exon 5 inclusion (**K&L**), SEPTIN6 exon 10 inclusion (**M&N**), and STAG2 exon 31 inclusion (**O&P**) was performed. Values represent averages of three independent experiments ±SD *p<0.05; Student’s t-test compared to PSI and total mRNA for control cells.

**Sub. Figure 5.** K562 cells were treated with thymidine at a final concentration of 2 mM, for 18 h. Cells were washed, then after 9 h treated again with 2 mM thymidine. After 18 h cells were washed and then treated with media or the PP2A inhibitor OA at a final concentration of 20 nM for 12 h. Cells were collected for RT-PCR. RNA was extracted, converted to cDNA, and total mRNA relative to CycloA and hTBP reference genes is shown for ARPP19 (**A**), ENSA (**B**). RT-PCR analysis of the PSI relative to RT-PCR analysis of total mRNA of ECT2 exon 5 inclusion (**C&D**), SEPTIN6 exon 10 inclusion (**E&F**), and STAG2 exon 31 inclusion (**G&H**) was performed. Values represent averages of three independent experiments ±SD *p<0.05; Student’s t-test. **I-P.** HeLa cells were treated under the same conditions and were harvested for RT-PCR. RNA was extracted, converted to cDNA and RT-PCR performed. Total mRNA relative to CycloA and hTBP reference genes is shown for ARPP19 (**I**), ENSA (**J**). RT-PCR analysis of the PSI relative to RT-PCR analysis of total mRNA of ECT2 exon 5 inclusion (**K&L**), SEPTIN6 exon 10 inclusion (**M&N**), and STAG2 exon 31 inclusion (**O&P**) was performed. Values represent averages of three independent experiments ±SD *p<0.05; Student’s t-test.

**Sup. Figure 6. A**-**B**. Kaplan-Meier curves for ECT2 exon 5 inclusion (**A**); and total mRNA of ECT2 (**B**). **C-D.** Kaplan-Meier curves for STAG2 exon 31 inclusion (**C**); and total mRNA of STAG2 (**D**). **E-F.** Kaplan-Meier curves for SEPTIN6 exon 10 inclusion (**E**); and total mRNA of SEPTIN6 (**F**). PSI>0 (red) and PSI=0 (blue). Error bars represent 95% confidence intervals of the mean.

**Sup. Figure 7. A-C.** ENSA-short/long were cloned to pLentic CMV GFP vector and lentiviral particles were produced by expressing ENSA short or long isoform in the HEK293T packaging cell-line using TransIT-X2 transfection reagent. HeLa cells were infected with viral particles to overexpress either the short or long isoform of ENSA. Protein was collected, and immunoblotting was conducted with the indicated antibodies and quantified relative to the housekeeping protein tubulin (**A**). The stable cell lines were arrested at G2/M using NCO for 20 h, then the cells were released for 3 h, collected and analyzed by flow cytometry (**B&C**).

## References

1. Wang, C., et al., Phosphorylation of spliceosomal protein SAP 155 coupled with splicing catalysis. Genes Dev, 1998. 12(10): p. 1409–14.

2. Girard, C., et al., Post-transcriptional spliceosomes are retained in nuclear speckles until splicing completion. Nat Commun, 2012. 3: p. 994.

3. Seghezzi, W., et al., Cyclin E associates with components of the pre-mRNA splicing machinery in mammalian cells. Mol Cell Biol, 1998. 18(8): p. 4526–36.

4. de Graaf, K., et al., The protein kinase DYRK1A phosphorylates the splicing factor SF3b1/SAP155 at Thr434, a novel in vivo phosphorylation site. BMC Biochem, 2006. 7: p. 7.

5. Boudrez, A., et al., Phosphorylation-dependent interaction between the splicing factors SAP155 and NIPP1. J Biol Chem, 2002. 277(35): p. 31834–41.

6. Tanuma, N., et al., Nuclear inhibitor of protein phosphatase-1 (NIPP1) directs protein phosphatase-1 (PP1) to dephosphorylate the U2 small nuclear ribonucleoprotein particle (snRNP) component, spliceosome-associated protein 155 (Sap155). J Biol Chem, 2008. 283(51): p. 35805–14.

7. Murthy, T., et al., Cyclin-dependent kinase 1 (CDK1) and CDK2 have opposing roles in regulating interactions of splicing factor 3B1 with chromatin. J Biol Chem, 2018. 293(26): p. 10220–10234.

8. Petrone, A., et al., Identification of Candidate Cyclin-dependent kinase 1 (Cdk1) Substrates in Mitosis by Quantitative Phosphoproteomics. Mol Cell Proteomics, 2016. 15(7): p. 2448–61.

9. Zhang, Y., et al., Inhibition of Splicing Factor 3b Subunit 1 (SF3B1) Reduced Cell Proliferation, Induced Apoptosis and Resulted in Cell Cycle Arrest by Regulating Homeobox A10 (HOXA10) Splicing in AGS and MKN28 Human Gastric Cancer Cells. Med Sci Monit, 2020. 26: p. e919460.

10. Popli, P., et al., Splicing factor SF3B1 promotes endometrial cancer progression via regulating KSR2 RNA maturation. Cell Death Dis, 2020. 11(10): p. 842.

11. Han, C., et al., SF3B1 homeostasis is critical for survival and therapeutic response in T cell leukemia. Sci Adv, 2022. 8(3): p. eabj8357.

12. Vanzyl, E.J., et al., Flow cytometric analysis identifies changes in S and M phases as novel cell cycle alterations induced by the splicing inhibitor isoginkgetin. PLoS One, 2018. 13(1): p. e0191178.

13. Salton, M. and T. Misteli, Small Molecule Modulators of Pre-mRNA Splicing in Cancer Therapy. Trends Mol Med, 2016. 22(1): p. 28–37.

14. Huang, Y., et al., SF3B1 deficiency impairs human erythropoiesis via activation of p53 pathway: implications for understanding of ineffective erythropoiesis in MDS. J Hematol Oncol, 2018. 11(1): p. 19.

15. Paolella, B.R., et al., Copy-number and gene dependency analysis reveals partial copy loss of wild-type SF3B1 as a novel cancer vulnerability. Elife, 2017. 6.

16. Pellagatti, A. and J. Boultwood, SF3B1 mutant myelodysplastic syndrome: Recent advances. Adv Biol Regul, 2021. 79: p. 100776.

17. Zhou, Z., et al., The biological function and clinical significance of SF3B1 mutations in cancer. Biomark Res, 2020. 8: p. 38.

18. Misteli, T., RNA splicing: What has phosphorylation got to do with it? Curr Biol, 1999. 9(6): p. R198–200.

19. Obeng, E.A., et al., Physiologic Expression of Sf3b1(K700E) Causes Impaired Erythropoiesis, Aberrant Splicing, and Sensitivity to Therapeutic Spliceosome Modulation. Cancer Cell, 2016. 30(3): p. 404–417.

20. Seiler, M., et al., H3B-8800, an orally available small-molecule splicing modulator, induces lethality in spliceosome-mutant cancers. Nat Med, 2018. 24(4): p. 497–504.

21. Lopez-Oreja, I., et al., SF3B1 mutation-mediated sensitization to H3B-8800 splicing inhibitor in chronic lymphocytic leukemia. Life Sci Alliance, 2023. 6(11).

22. Petasny, M., et al., Splicing to Keep Cycling: The Importance of Pre-mRNA Splicing during the Cell Cycle. Trends Genet, 2021. 37(3): p. 266–278.

23. Love, M.I., W. Huber, and S. Anders, Moderated estimation of fold change and dispersion for RNA-seq data with DESeq2. Genome Biol, 2014. 15(12): p. 550.

24. Shen, S., et al., rMATS: robust and flexible detection of differential alternative splicing from replicate RNA-Seq data. Proc Natl Acad Sci U S A, 2014. 111(51): p. E5593–601.

25. Ben-Ari Fuchs, S., et al., GeneAnalytics: An Integrative Gene Set Analysis Tool for Next Generation Sequencing, RNAseq and Microarray Data. OMICS, 2016. 20(3): p. 139–51.

26. Zhang, Y., et al., OncoSplicing: an updated database for clinically relevant alternative splicing in 33 human cancers. Nucleic Acids Res, 2022. 50(D1): p. D1340–D1347.

27. Wang, L., et al., Transcriptomic Characterization of SF3B1 Mutation Reveals Its Pleiotropic Effects in Chronic Lymphocytic Leukemia. Cancer Cell, 2016. 30(5): p. 750–763.

28. Zhang, Y., et al., OncoSplicing 3.0: an updated database for identifying RBPs regulating alternative splicing events in cancers. Nucleic Acids Res, 2025. 53(D1): p. D1460–D1466.

29. Tang, Z., et al., GEPIA: a web server for cancer and normal gene expression profiling and interactive analyses. Nucleic Acids Res, 2017. 45(W1): p. W98–W102.

30. Padi, S.K.R., et al., Cryo-EM structures of PP2A:B55-FAM122A and PP2A:B55-ARPP19. Nature, 2024. 625(7993): p. 195–203.

