## Supplementary material for "SF3B1^K700E^ rewires splicing of cell-cycle regulators": sup fig

Sup. Fig. 1

**A**

| Event Type | All Significant Events | Significant Events WT Higher Inclusion | Significant Events SF3B1 MUT Higher Inclusion |
|---|---|---|---|
| **SE** | 8953 | 4437 | 4516 |
| **A5SS** | 1098 | 510 | 588 |
| **A3SS** | 3583 | 1531 | 2052 |
| **MXE** | 7408 | 3480 | 3928 |
| **RI** | 1094 | 664 | 430 |

**B**

**Upregulated genes**

| **Pathway** | **Significance (-Log10(P-value))** | **# Matched genes** |
|---|---|---|
| Akt signaling | 20.3 | 34 |
| **Phospholipase-C Pathway** | **20.03** | **20** |
| Angiogenesis | 19 | 11 |
| **Interleukin-10 Signaling** | **17.61** | **[illegible]** |
| ECM Proteoglycans | 17.32 | 12 |

**Downregulated genes**

| **Pathway** | **Significance (-Log10(P-value))** | **# Matched genes** |
|---|---|---|
| Influenza Infection | 15.6 | 8 |
| **Metal Ion SLC Transporters** | **15.27** | **[illegible]** |
| Mitochondrial Unfolded Protein Response (UPRmt) | 11.63 | 3 |
| **Copper Homeostasis** | **[illegible]** | **4** |
| Antiviral Mechanism By IFN-stimulated Genes | 10.35 | 8 |

**C**

**ARPP19**
chr15: 52856387-52856443

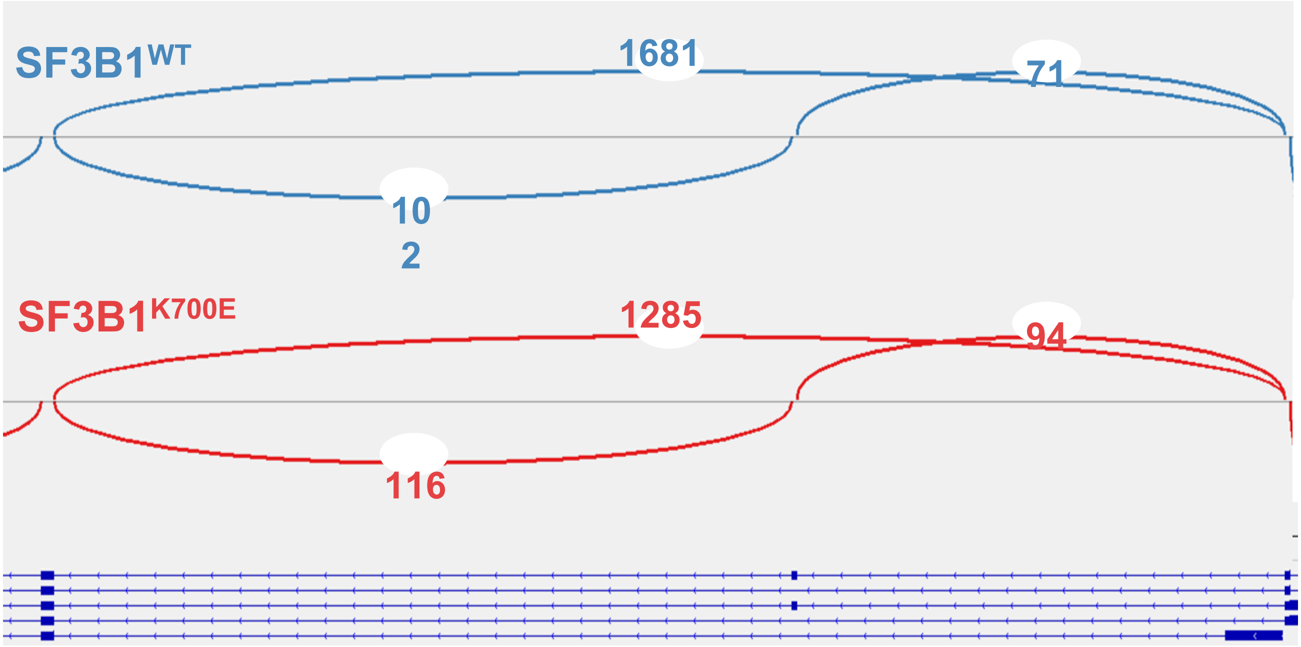

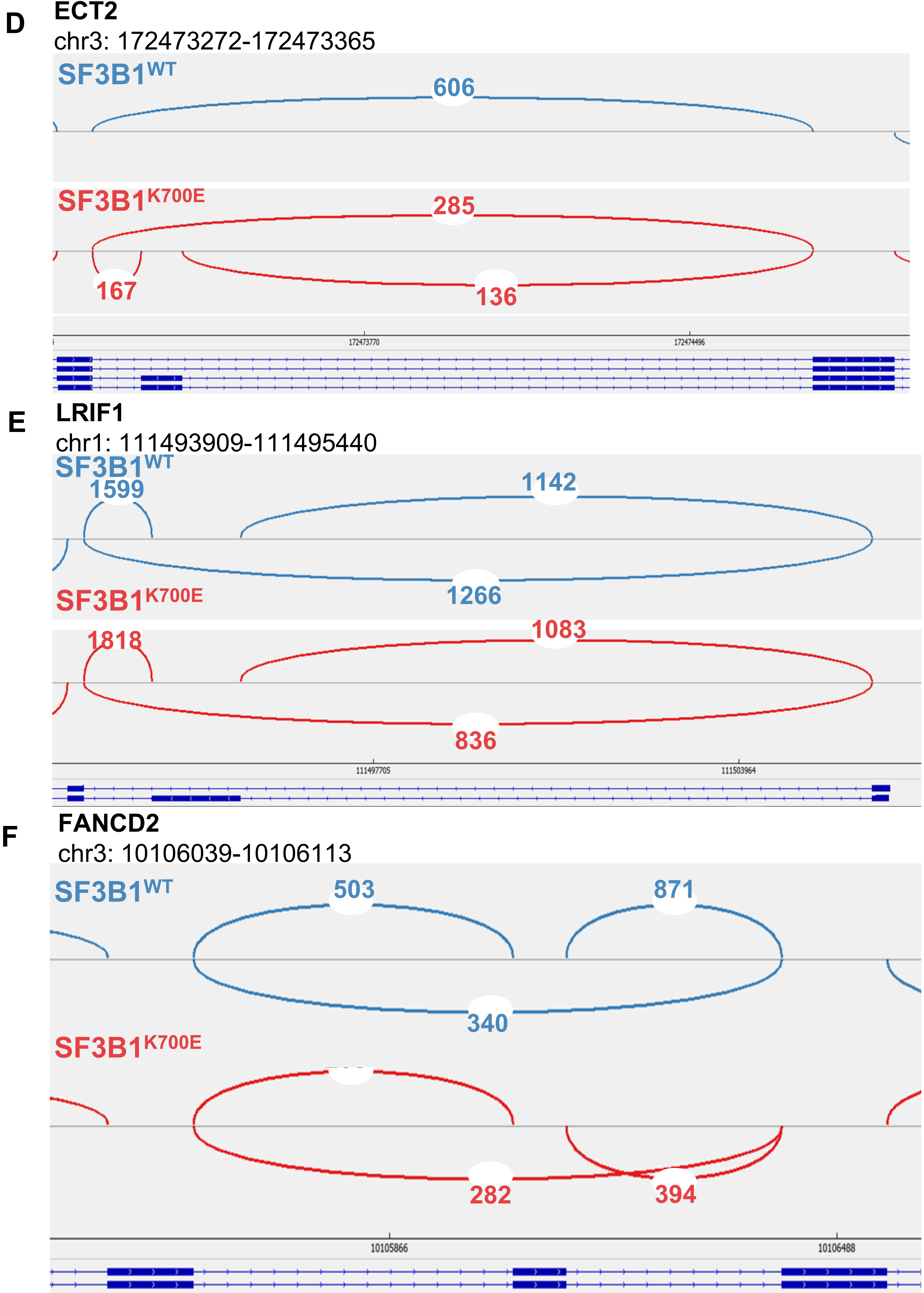
D
ECT2
chr3: 172473272-172473365
SF3B1WT
606
SF3B1K700E
285
167
136
172473770
172474496
E
LRIF1
chr1: 111493909-111495440
SF3B1WT
1599
1142
1266
SF3B1K700E
1818
1083
836
111497705
111503964
F
FANCD2
chr3: 10106039-10106113
SF3B1WT
503
871
340
SF3B1K700E
282
394
10105866
10106488

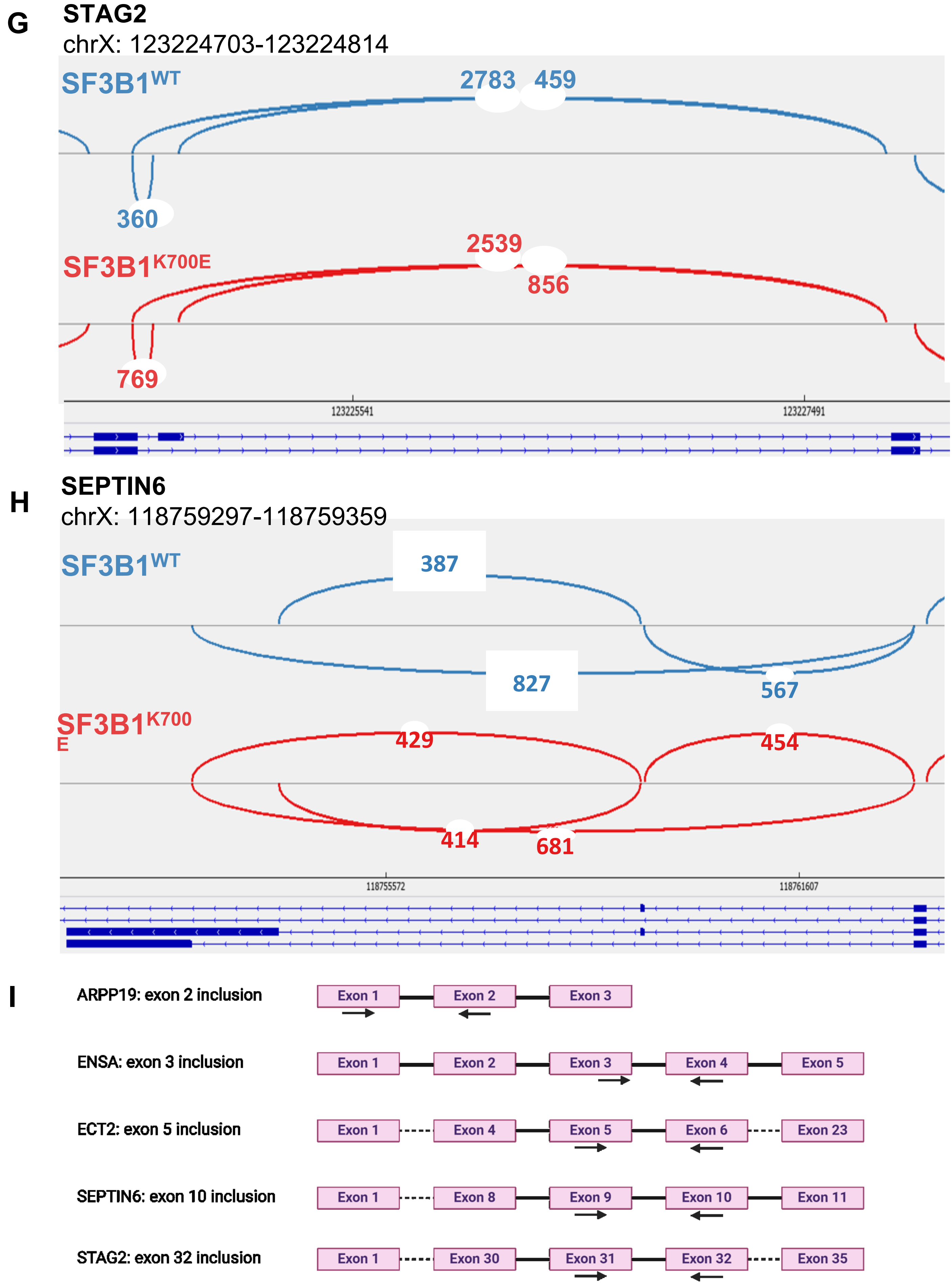
G
STAG2
chrX: 123224703-123224814
SF3B1WT
2783
459
360
SF3B1K700E
2539
856
769
123225541
123227491
H
SEPTIN6
chrX: 118759297-118759359
SF3B1WT
387
827
567
SF3B1K700E
429
454
414
681
118755572
118761607
I
ARPP19: exon 2 inclusion
Exon 1
Exon 2
Exon 3
ENSA: exon 3 inclusion
Exon 1
Exon 2
Exon 3
Exon 4
Exon 5
ECT2: exon 5 inclusion
Exon 1
Exon 4
Exon 5
Exon 6
Exon 23
SEPTIN6: exon 10 inclusion
Exon 1
Exon 8
Exon 9
Exon 10
Exon 11
STAG2: exon 32 inclusion
Exon 1
Exon 30
Exon 31
Exon 32
Exon 35

Sup. Fig. 2

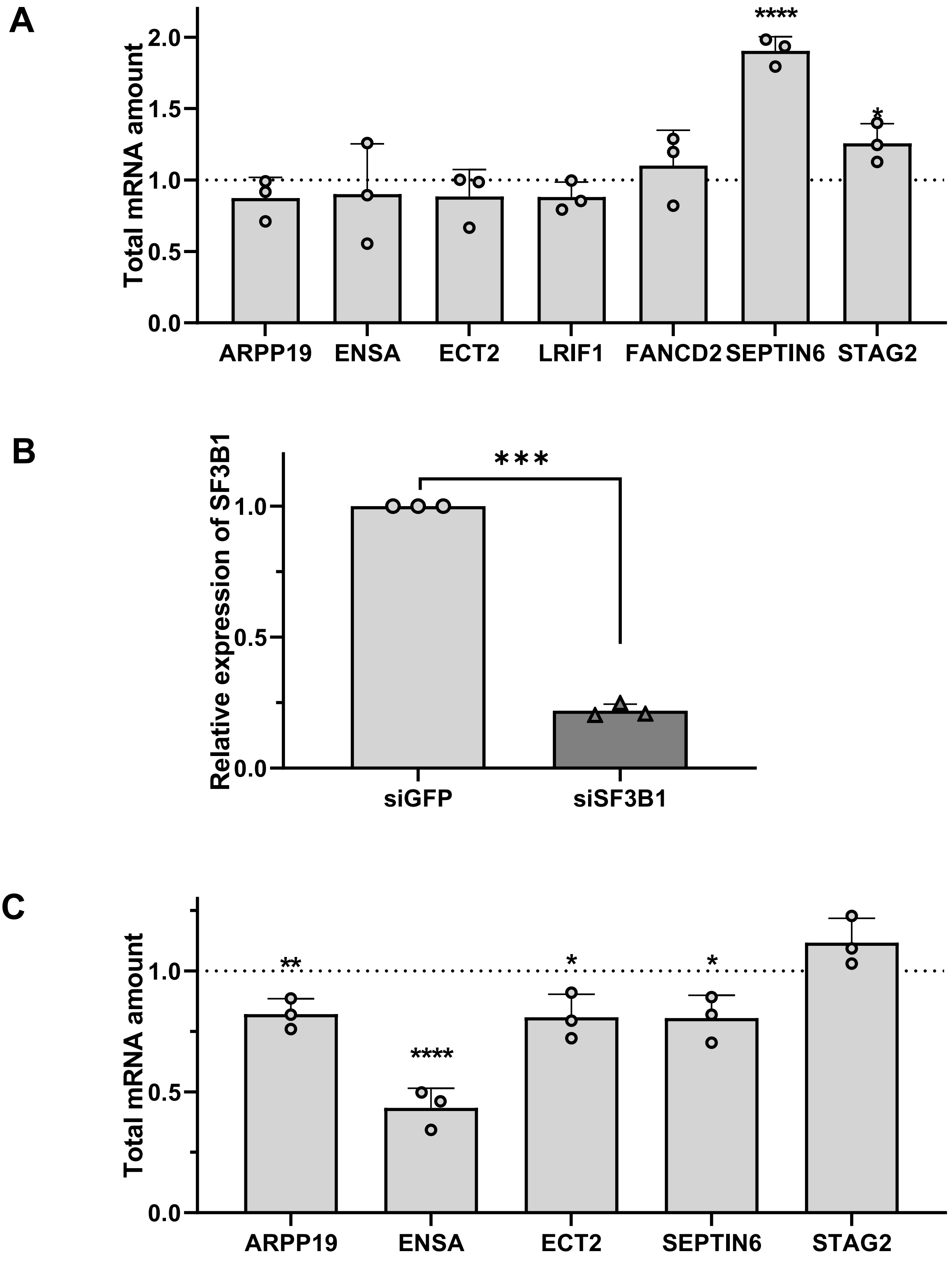
A
Total mRNA amount
2.0
1.5
1.0
0.5
0.0
****
*
ARPP19
ENSA
ECT2
LRIF1
FANCD2
SEPTIN6
STAG2
B
Relative expression of SF3B1
***
1.0
0.5
0.0
siGFP
siSF3B1
C
Total mRNA amount
1.0
0.5
0.0
**
****
*
*
ARPP19
ENSA
ECT2
SEPTIN6
STAG2

Sup. Fig. 3

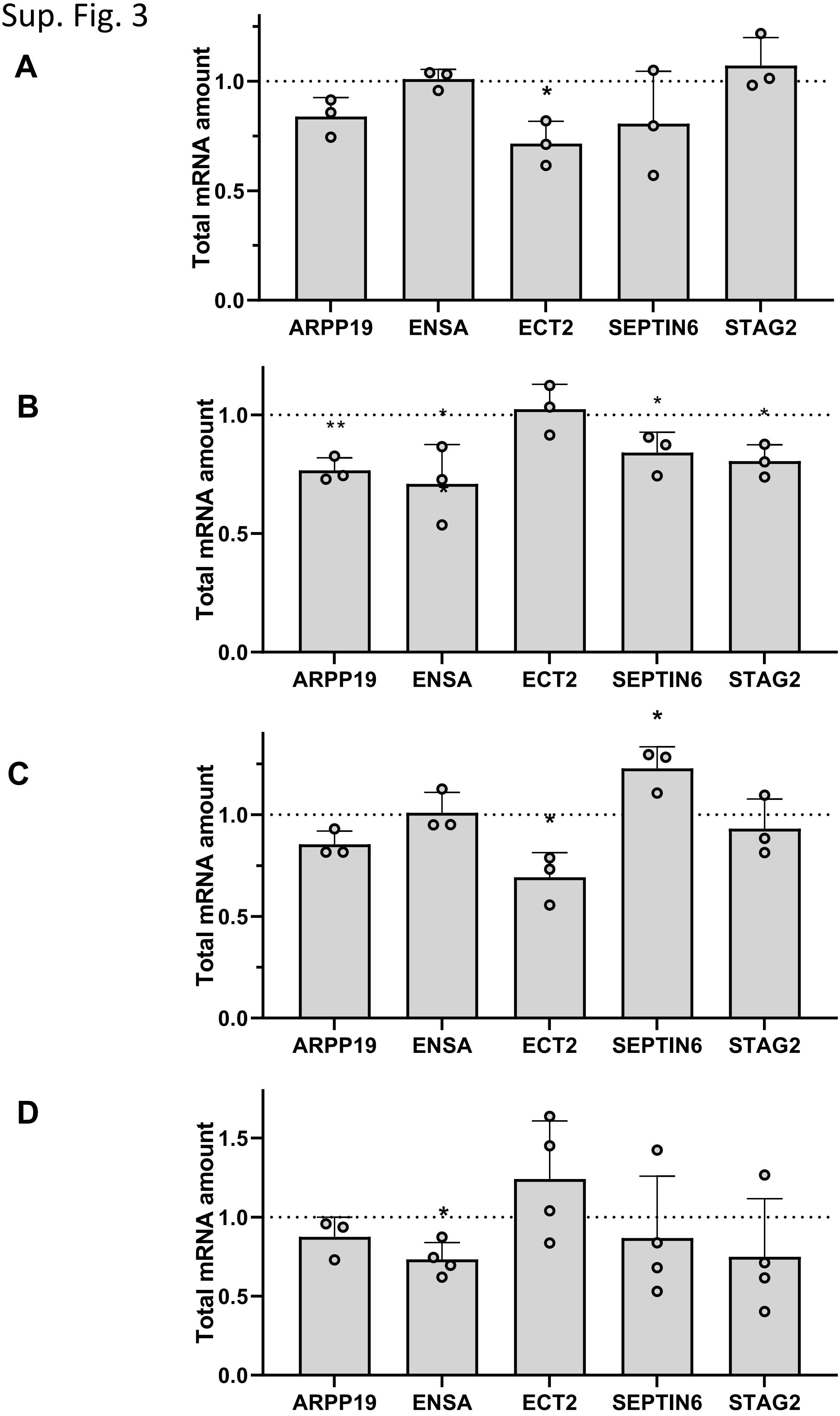
A
B
C
D
Total mRNA amount
0.0
0.5
1.0
1.5
ARPP19
ENSA
ECT2
SEPTIN6
STAG2
*
**

Sup. Fig. 4

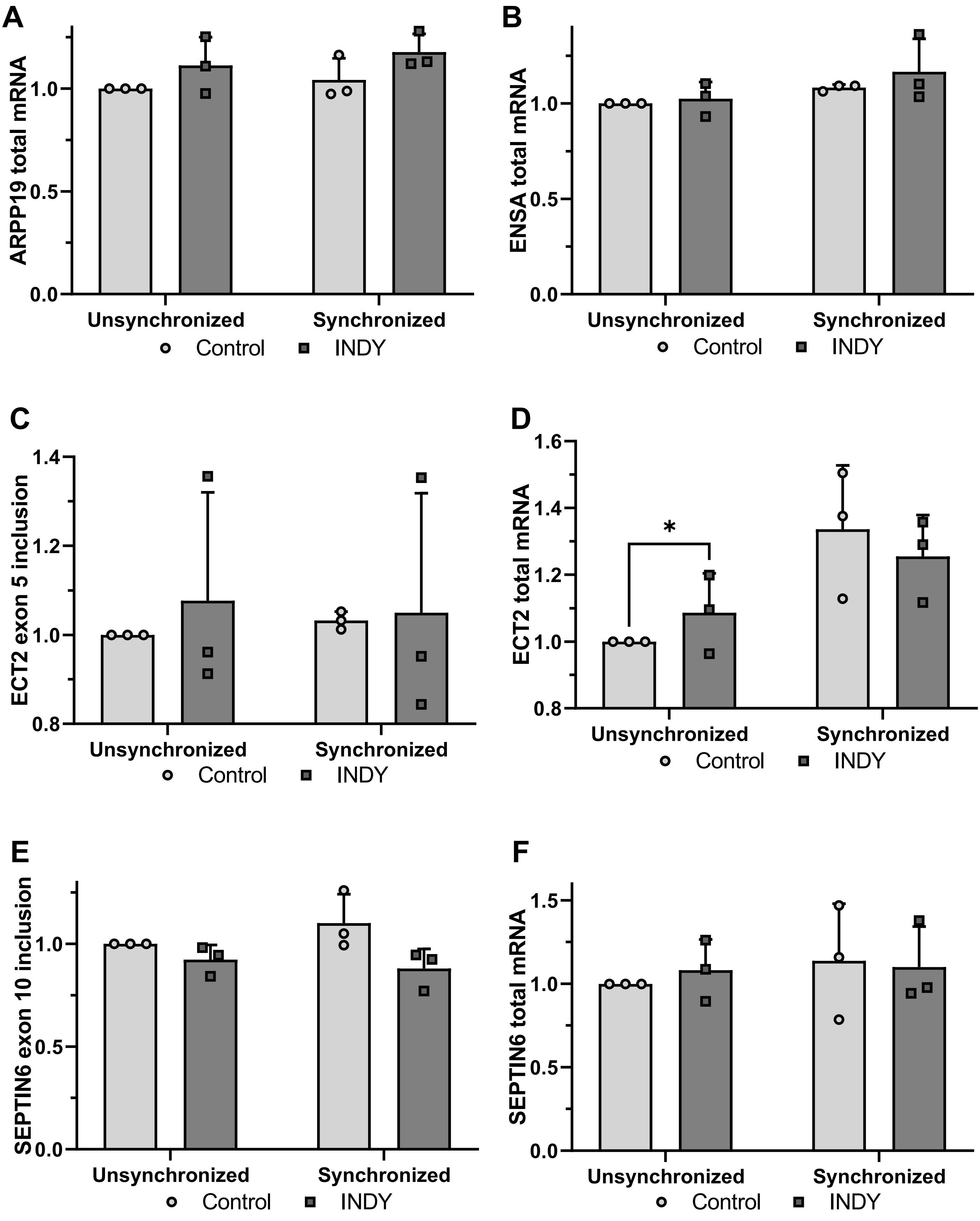
A
ARPP19 total mRNA
0.0
0.5
1.0
Unsynchronized
Synchronized
Control
INDY
B
ENSA total mRNA
0.0
0.5
1.0
Unsynchronized
Synchronized
Control
INDY
C
ECT2 exon 5 inclusion
0.8
1.0
1.2
1.4
Unsynchronized
Synchronized
Control
INDY
D
ECT2 total mRNA
0.8
1.0
1.2
1.4
1.6
*
Unsynchronized
Synchronized
Control
INDY
E
SEPTIN6 exon 10 inclusion
0.0
0.5
1.0
Unsynchronized
Synchronized
Control
INDY
F
SEPTIN6 total mRNA
0.0
0.5
1.0
1.5
Unsynchronized
Synchronized
Control
INDY

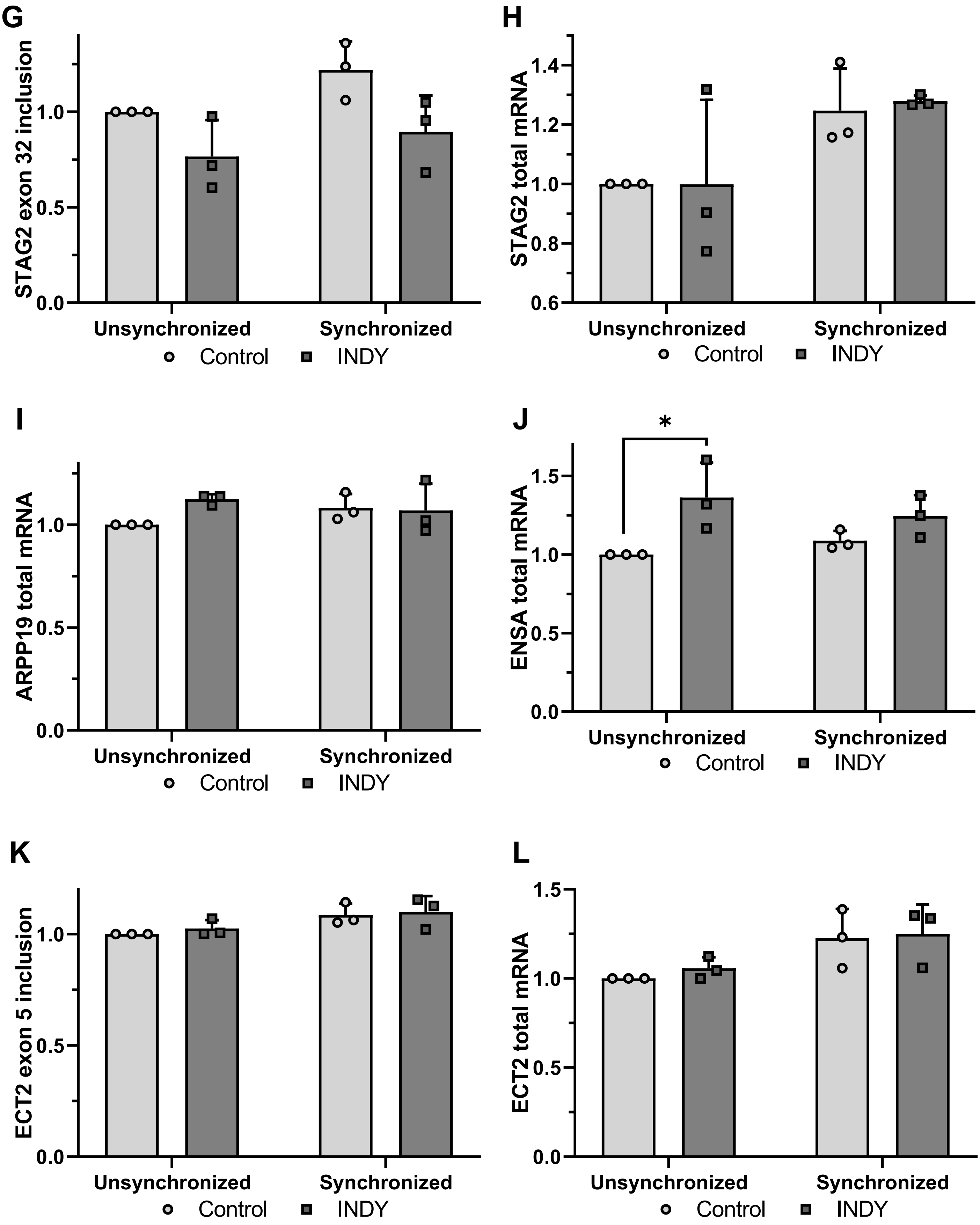

G
STAG2 exon 32 inclusion
0.0
0.5
1.0
Unsynchronized
Synchronized
Control
INDY
H
STAG2 total mRNA
0.6
0.8
1.0
1.2
1.4
Unsynchronized
Synchronized
Control
INDY
I
ARPP19 total mRNA
0.0
0.5
1.0
Unsynchronized
Synchronized
Control
INDY
J
*
ENSA total mRNA
0.0
0.5
1.0
1.5
Unsynchronized
Synchronized
Control
INDY
K
ECT2 exon 5 inclusion
0.0
0.5
1.0
Unsynchronized
Synchronized
Control
INDY
L
ECT2 total mRNA
0.0
0.5
1.0
1.5
Unsynchronized
Synchronized
Control
INDY

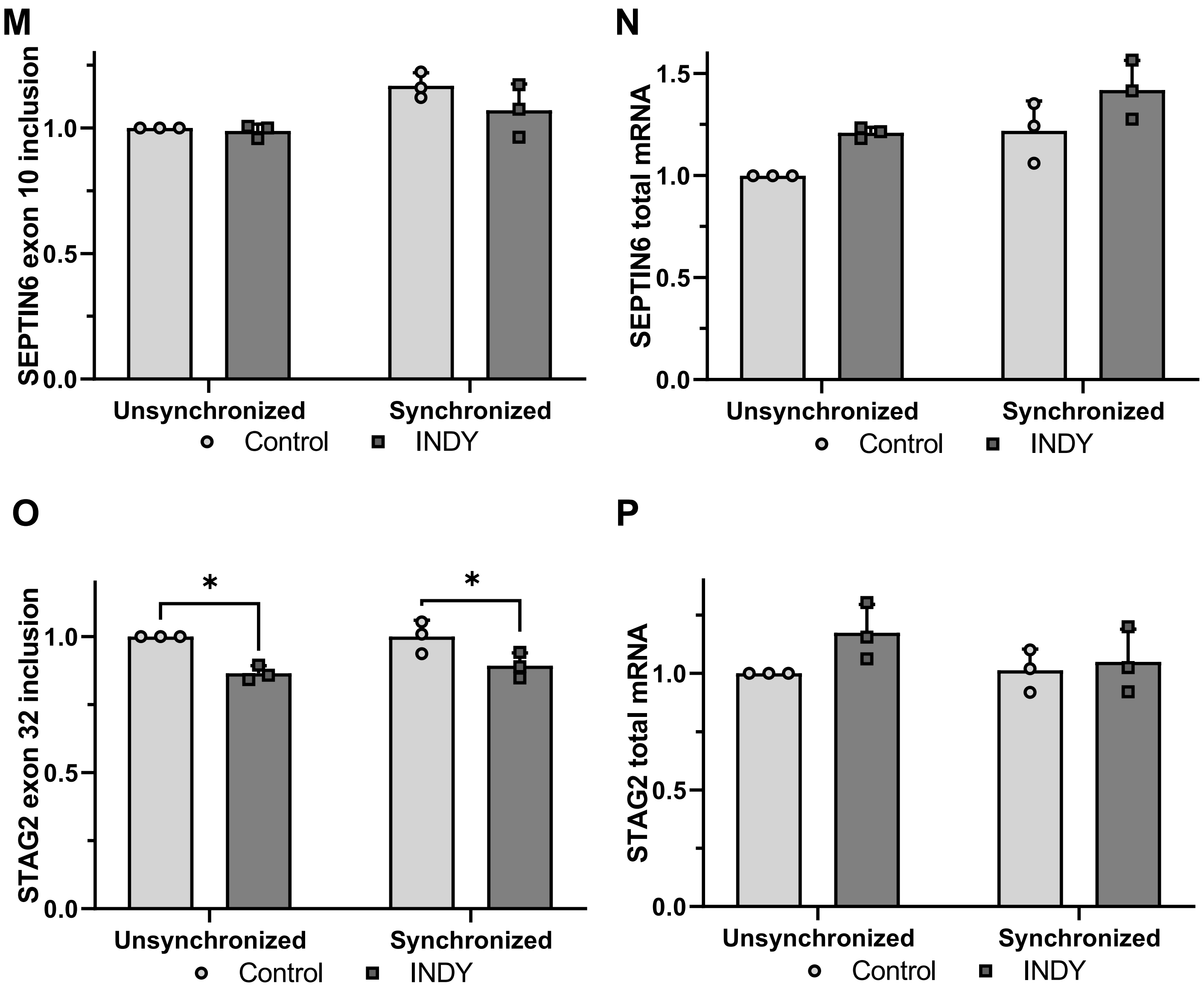
M
SEPTIN6 exon 10 inclusion
0.0
0.5
1.0
Unsynchronized
Synchronized
Control
INDY
N
SEPTIN6 total mRNA
0.0
0.5
1.0
1.5
Unsynchronized
Synchronized
Control
INDY
O
STAG2 exon 32 inclusion
*
*
0.0
0.5
1.0
Unsynchronized
Synchronized
Control
INDY
P
STAG2 total mRNA
0.0
0.5
1.0
Unsynchronized
Synchronized
Control
INDY

Sup. Fig. 5

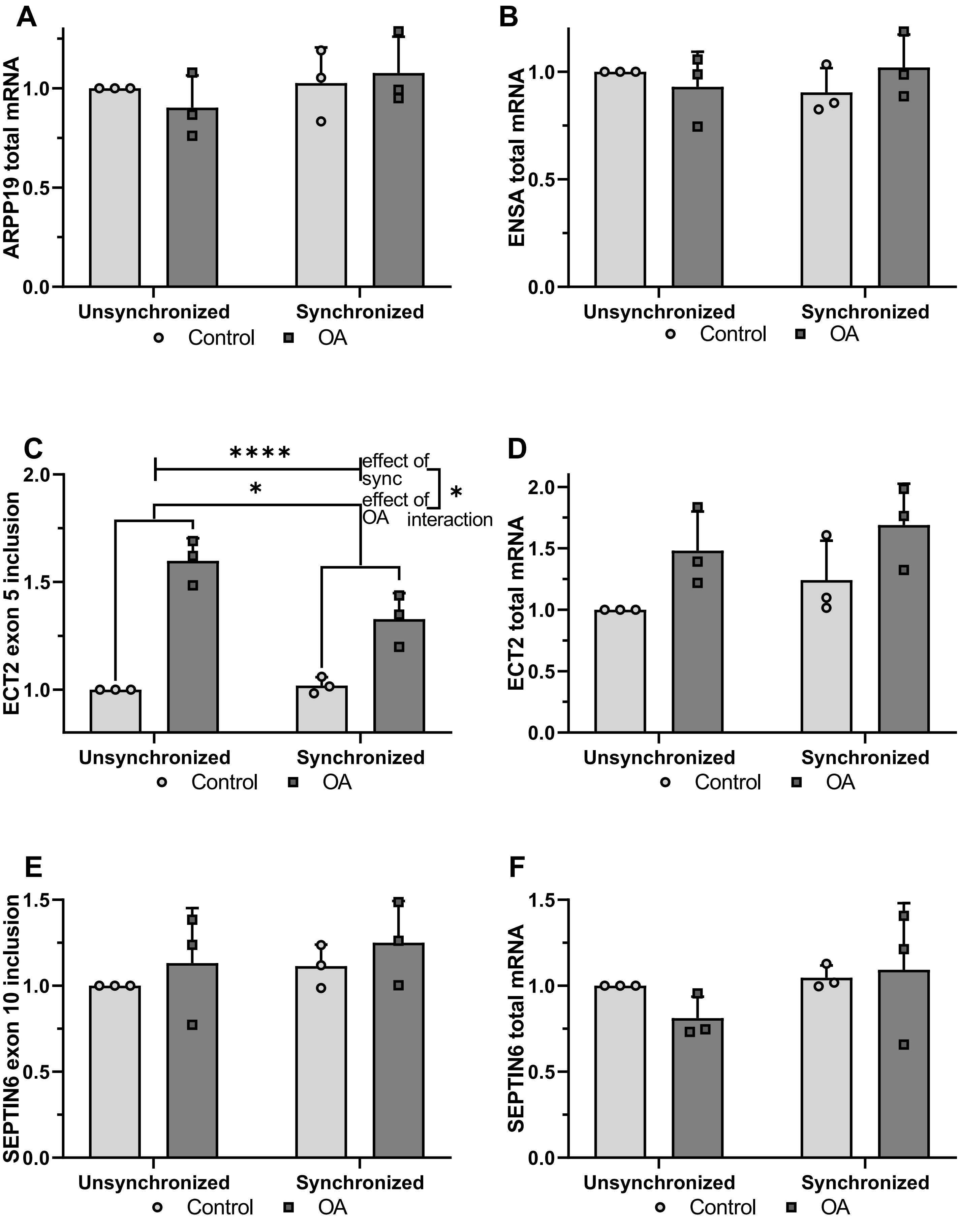
A
ARPP19 total mRNA
0.0
0.5
1.0
Unsynchronized
Synchronized
Control
OA
B
ENSA total mRNA
0.0
0.5
1.0
Unsynchronized
Synchronized
Control
OA
C
ECT2 exon 5 inclusion
1.0
1.5
2.0
****
*
effect of sync
effect of OA
*
interaction
Unsynchronized
Synchronized
Control
OA
D
ECT2 total mRNA
0.0
0.5
1.0
1.5
2.0
Unsynchronized
Synchronized
Control
OA
E
SEPTIN6 exon 10 inclusion
0.0
0.5
1.0
1.5
Unsynchronized
Synchronized
Control
OA
F
SEPTIN6 total mRNA
0.0
0.5
1.0
1.5
Unsynchronized
Synchronized
Control
OA

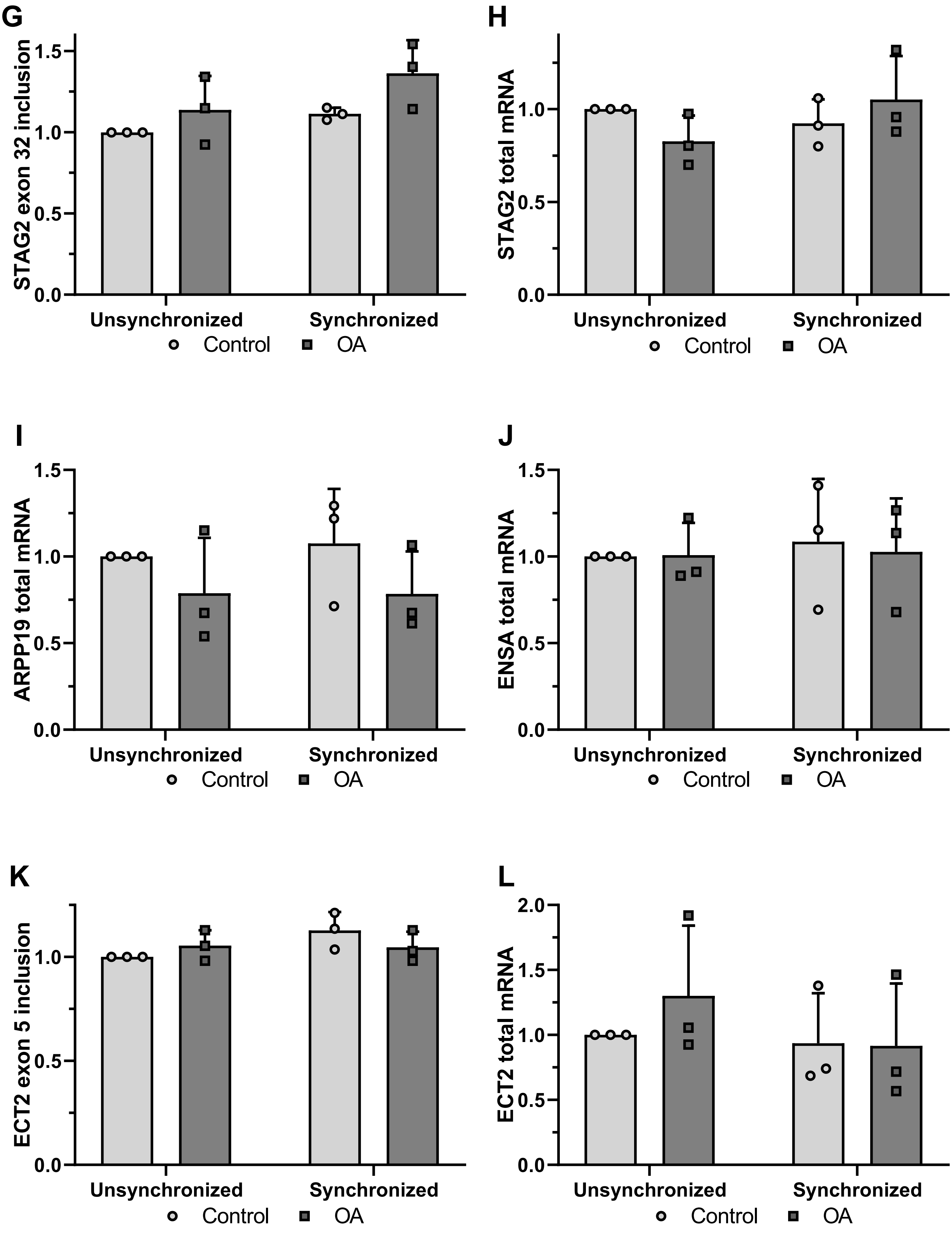
G
STAG2 exon 32 inclusion
1.5
1.0
0.5
0.0
Unsynchronized
Synchronized
Control
OA
H
STAG2 total mRNA
1.0
0.5
0.0
Unsynchronized
Synchronized
Control
OA
I
ARPP19 total mRNA
1.5
1.0
0.5
0.0
Unsynchronized
Synchronized
Control
OA
J
ENSA total mRNA
1.5
1.0
0.5
0.0
Unsynchronized
Synchronized
Control
OA
K
ECT2 exon 5 inclusion
1.0
0.5
0.0
Unsynchronized
Synchronized
Control
OA
L
ECT2 total mRNA
2.0
1.5
1.0
0.5
0.0
Unsynchronized
Synchronized
Control
OA

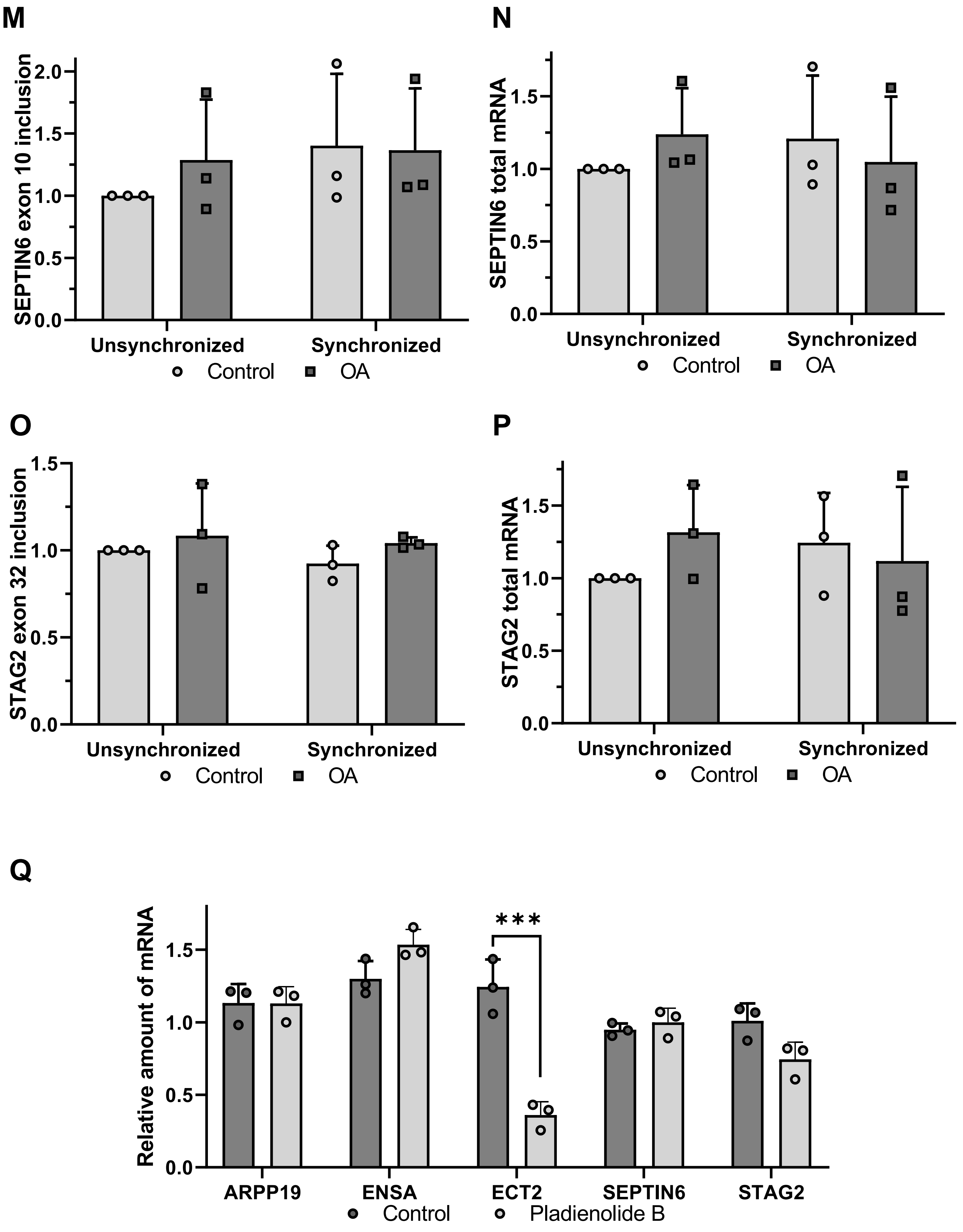
M
SEPTIN6 exon 10 inclusion
0.0
0.5
1.0
1.5
2.0
Unsynchronized
Synchronized
Control
OA
N
SEPTIN6 total mRNA
0.0
0.5
1.0
1.5
Unsynchronized
Synchronized
Control
OA
O
STAG2 exon 32 inclusion
0.0
0.5
1.0
1.5
Unsynchronized
Synchronized
Control
OA
P
STAG2 total mRNA
0.0
0.5
1.0
1.5
Unsynchronized
Synchronized
Control
OA
Q
Relative amount of mRNA
0.0
0.5
1.0
1.5
***
ARPP19
ENSA
ECT2
SEPTIN6
STAG2
Control
Pladienolide B

Sup. Fig. 6

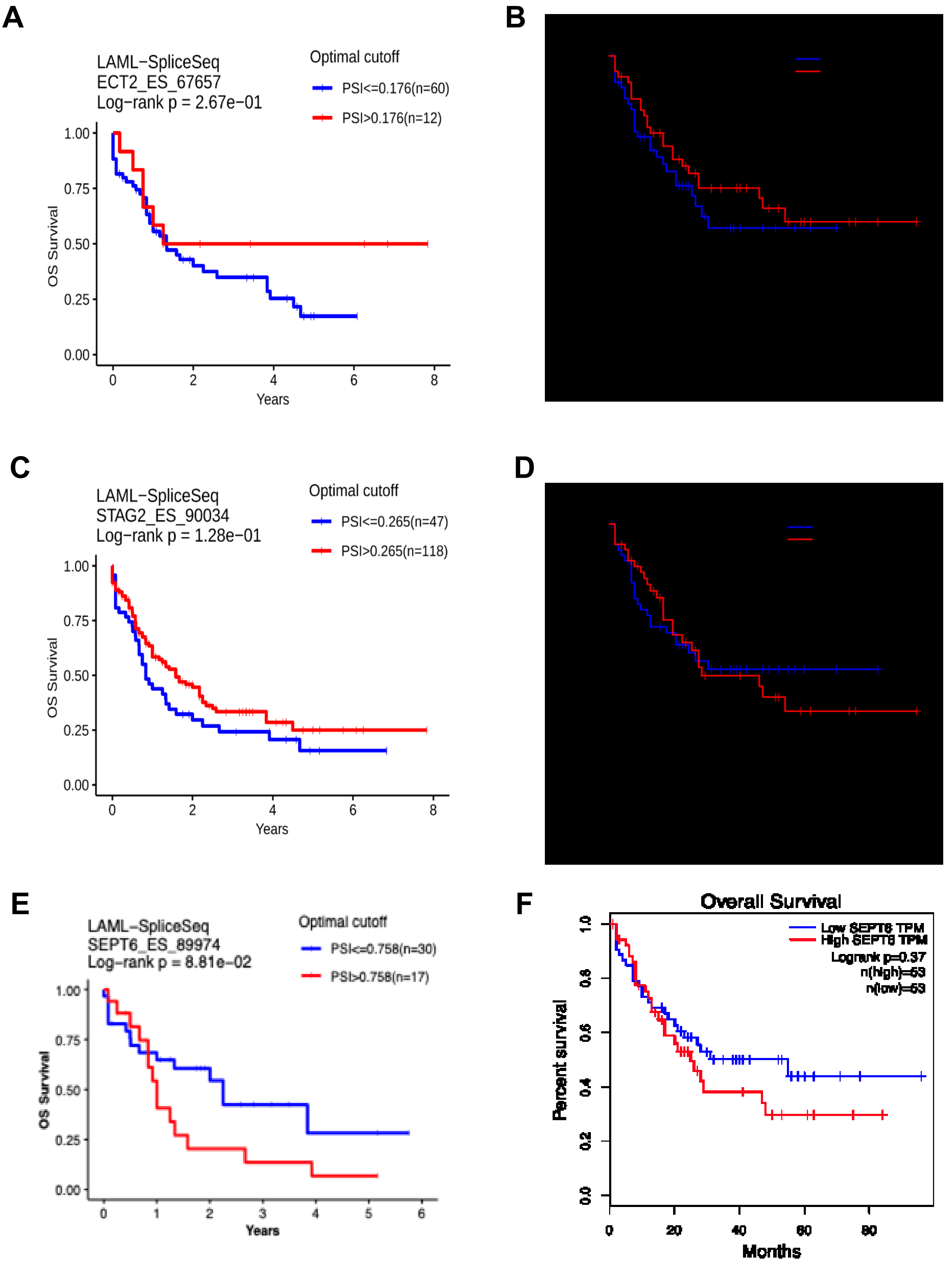
A
LAML−SpliceSeq
ECT2_ES_67657
Log−rank p = 2.67e−01
Optimal cutoff
PSI<=0.176(n=60)
PSI>0.176(n=12)
OS Survival
1.00
0.75
0.50
0.25
0.00
0
2
4
6
8
Years
B
C
LAML−SpliceSeq
STAG2_ES_90034
Log−rank p = 1.28e−01
Optimal cutoff
PSI<=0.265(n=47)
PSI>0.265(n=118)
OS Survival
1.00
0.75
0.50
0.25
0.00
0
2
4
6
8
Years
D
E
LAML−SpliceSeq
SEPT6_ES_89974
Log−rank p = 8.81e−02
Optimal cutoff
PSI<=0.758(n=30)
PSI>0.758(n=17)
OS Survival
1.00
0.75
0.50
0.25
0.00
0
1
2
3
4
5
6
Years
F
Overall Survival
Low SEPT6 TPM
High SEPT6 TPM
Logrank p=0.37
n(high)=53
n(low)=53
Percent survival
1.0
0.8
0.6
0.4
0.2
0.0
0
20
40
60
80
Months

Sup. Fig. 7

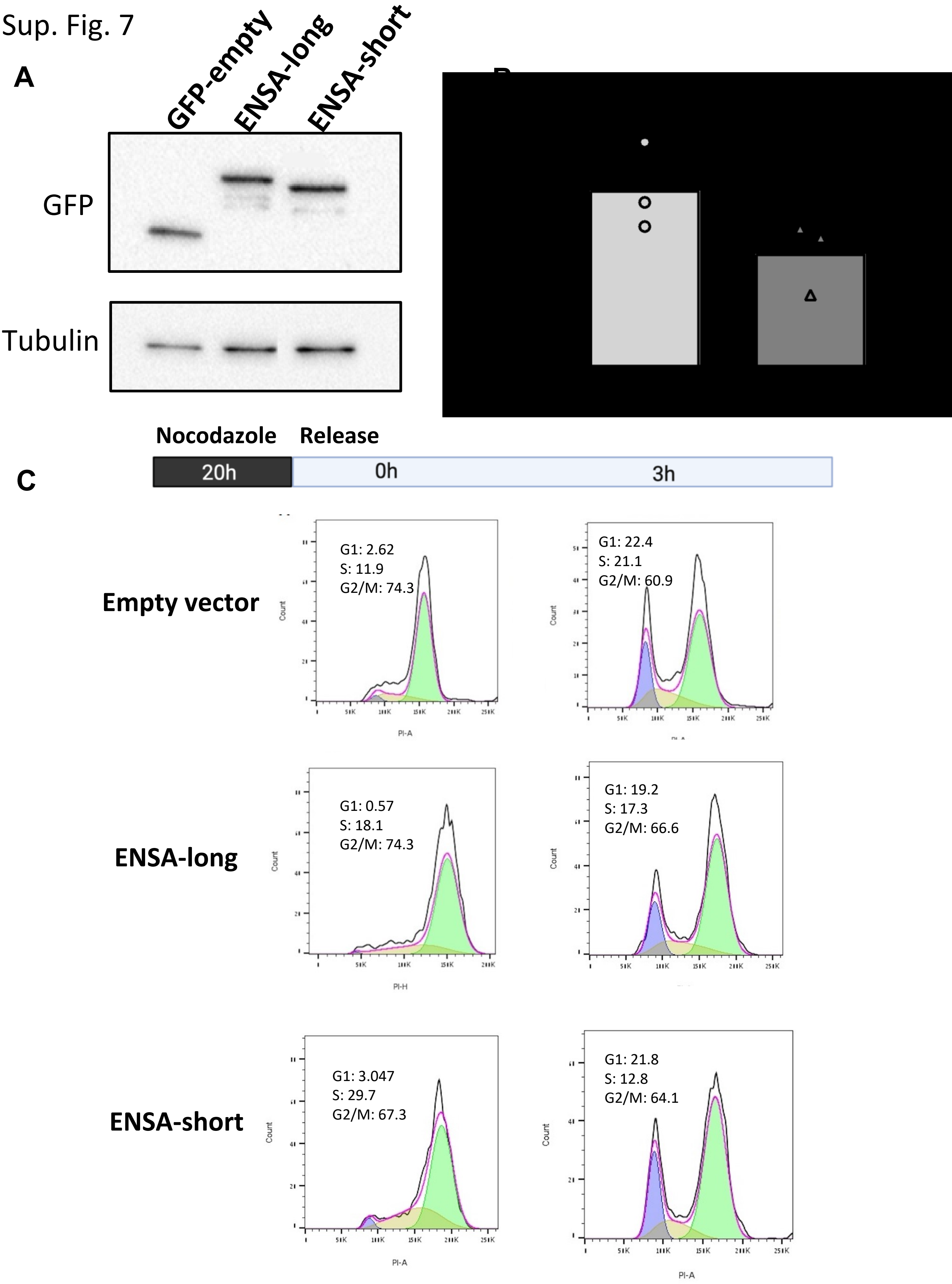
A
GFP-empty
ENSA-long
ENSA-short
GFP
Tubulin
C
Nocodazole
Release
20h
0h
3h
Empty vector
G1: 2.62
S: 11.9
G2/M: 74.3
G1: 22.4
S: 21.1
G2/M: 60.9
ENSA-long
G1: 0.57
S: 18.1
G2/M: 74.3
G1: 19.2
S: 17.3
G2/M: 66.6
ENSA-short
G1: 3.047
S: 29.7
G2/M: 67.3
G1: 21.8
S: 12.8
G2/M: 64.1
Count
PI-A
PI-H
